# PD-1 dependent expansion of Amphregulin^+^FOXP3^+^ cells is associated with oral immune dysfunction in HIV patients on therapy

**DOI:** 10.1101/2021.03.13.435273

**Authors:** N Bhaskaran, E Schneider, F Faddoul, A Paes da Silva, R Asaad, A Talla, N Greenspan, AD Levine, D McDonald, J Karn, MM Lederman, P Pandiyan

## Abstract

Residual systemic inflammation and mucosal immune dysfunction persist in people living with HIV (PLWH) despite treatment with combined anti-retroviral therapy (cART), but the underlying immune mechanisms are poorly understood. Here we report an altered immune landscape involving upregulation of TLR- and inflammasome signaling, localized CD4^+^ T cell hyperactivation, and counterintuitively, an enrichment of CD4^+^CD25^+^FOXP3^+^ regulatory T cells (T_regs_) in the oral mucosa of HIV^+^ patients on therapy. Using human oral tonsil cultures, we found that HIV infection causes an increase in a unique population of FOXP3^+^ cells expressing PD-1, IFN-γ, Amphiregulin (AREG), and IL-10. These cells persisted even in the presence of the anti-retroviral drug and underwent further expansion driven by TLR-2 ligands and IL-1β. IL-1β also promoted PD-1 upregulation in AKT1 dependent manner. PD-1 stabilized FOXP3 and AREG expression in these cells through a mechanism requiring the activation of Asparaginyl Endopeptidase (AEP). Importantly, these FOXP3^+^ cells were incapable of suppressing CD4^+^ T cells *in vitro*. Concurrently, HIV^+^ patients harbored higher levels of PD-1, IFN-γ, Amphiregulin (AREG), and IL-10 expressing FOXP3^+^ cells, which strongly correlated with CD4^+^ T cell hyperactivation, suggesting an absence of CD4^+^ T cell regulation in the oral mucosa. Taken together, this study provides insights into a novel mechanism of FOXP3^+^ cell dysregulation and reveals a critical link in the positive feedback loop of oral mucosal immune activation events in HIV^+^ patients on therapy.

**One Sentence Summary:** HIV-induced immune dysfunction in lymphoid and mucosal tissues

## Introduction

Human immunodeficiency virus 1 (HIV-1) associated co-morbidities such as inflammatory disorders and cancer are important public health concerns^1–6^. Immune complications persist in patients despite effective combined antiretroviral therapy (cART), and have been inextricably linked to HIV latency, altered mucosal T cell functionality, and increased production of immune activation-associated cytokines in treated HIV^+^patients^7–14^. Although oral complications such as periodontitis and oropharyngeal cancer in healthy HIV-uninfected adults are usually mild, self-limited, and of short duration, they are of increased incidence and severity in HIV^+^ individuals under suppressive HIV therapy ^15–18^. Oral mucosa is conferred with a distinct immune compartment with a unique microbiome^19^, but the oral lymphoid cell population and its dysregulation in HIV^+^ patients are not understood ^18, 20, 21^. Acute simian immunodeficiency virus (SIV) infection has been shown to cause a loss of barrier protection as a result of CD4^+^ T cell depletion in oral mucosa^22^. Although cART therapy can restore these CD4^+^ T cells, they can contribute to oral viral reservoirs. To date, there is no information on alterations of oral mucosal CD4^+^ T cell functionality in the context of SIV or HIV infection after treatment. CD4^+^CD25^+^Foxp3^+^ T_reg_ cells, known for their immunomodulatory functions, express CXCR4 and CCR5 coreceptors and support high levels of HIV infection and replication^23^. Thus, the initial loss of T_regs_ during HIV infection can contribute to a self-perpetuating loop of events leading to immune activation^7, 24–28^. Previous reports document varied levels of T_regs_, depending on the location (blood, lymphoid organ, or mucosa) and acute *versus* chronic phase of infection^29–34^. Nevertheless, the precise cellular and functional alterations in T_regs_ in the context of immune activation have not been characterized in the oral mucosa. Here we show that gingival mucosa of cART-treated people living with HIV (PLWH) had an increased accrual of CD4^+^CD25^+^FOXP3^+^ cells when compared to healthy individuals. Counterintuitively, conventional CD4^+^ T cells showed a hyperactivated CD38^high^HLA-DR^+^ phenotype and not a “suppressed” phenotype in this environment. Transcriptomic profiling of the bulk immune population in gingival mucosa revealed an upregulation of TLR signaling and inflammasome pathway in PLWH. Examining FOXP3^+^ cells in HIV-infected oral lymphoid tonsillar cultures, we found that HIV, TLR-2 ligands, and IL-1β were capable of expanding a unique population of FOXP3^+^ cells expressing PD-1, IFN-γ, and AREG. These cells required IL-1β mediated AKT1 (AKT) phosphorylation and PD-1 dependent AEP activation for expansion and the expression of FOXP3 and AREG. However, these cells did not suppress CD4^+^ T cells *in vitro*, implying that T_reg_ mediated mucosal CD4^+^ T cell regulation could be impaired in HIV^+^ patients *in vivo*. Concurringly, the frequency of PD-1^hi^IFN-γ expressing FOXP3^+^ cells with an elevated expression of AREG and IL-10 was also higher in oral mucosa of HIV^+^ patients on therapy and correlated with the dysregulated immune landscape. Taken together, our results here provide significant mechanistic insights into HIV-mediated T_reg_ dysfunction in oral mucosa and unveil new targets to modulate immune activation.

## Results

### Oral gingival tissue displays inflammatory signature and CD4 T cell alterations in HIV^+^ patients on therapy

To examine immune cell alterations in oral mucosa, we recruited 78 participants that included healthy controls and treated HIV^+^ (HIV+ cART) individuals and collected their saliva, peripheral blood mononuclear cells (PBMC), and oral gingival mucosa biopsies (**Table 1**). Unbiased RNA sequencing analyses revealed an upregulation of 772 transcripts and downregulation of 226 transcripts in gingival biopsy tissues of HIV+ cART individuals when compared to controls (**Fig.1A, left**). However, only 54 genes were differentially regulated in their PBMC (**Fig.1A, right**), indicating a mucosal dysfunction persisting during therapy after significant clearance of the virus. Global pathway analysis identified that a majority of the upregulated genes in oral mucosa of HIV^+^ patients were associated with TLR, MyD88, inflammasome, and inflammatory responses, highlighting an underlying immune activation (**Fig.1B-D**). Gene set enrichment analysis (GSEA)^35^ revealed a positive enrichment of pathways of aging, head and neck cancer, and AKT1 signaling based on gene sets in GO pathways and MSigDB (**Fig.S1A**). The frequency of CD38 and HLA-DR co-expressing cells, the hallmark of HIV-mediated CD4^+^T cell activation^36^, was significantly higher in human oral intraepithelial and lamina propria leukocytes (HOIL) from gingival biopsies from HIV^+^ patients on therapy (**Fig.1E; Fig.S1B, S1C**). As we have shown previously, there were no differences in the frequency of activated CD4^+^ T cells between the PBMC of the groups (data not shown)^36^. Neither were there any differences in overall CD4^+^ T cell proportions or the levels of IFN-γ expressing CD4^+^ cells between these groups (**S2, A-C**). Collectively, peripheral CD4+ T cells appear to be largely restored by cART, but oral mucosa of HIV^+^ patients display features of immune dysregulation with alterations in inflammasome pathway, TLR/MyD88 signaling, and localized CD4^+^ T lymphocyte hyperactivation.

**Fig. 1:**
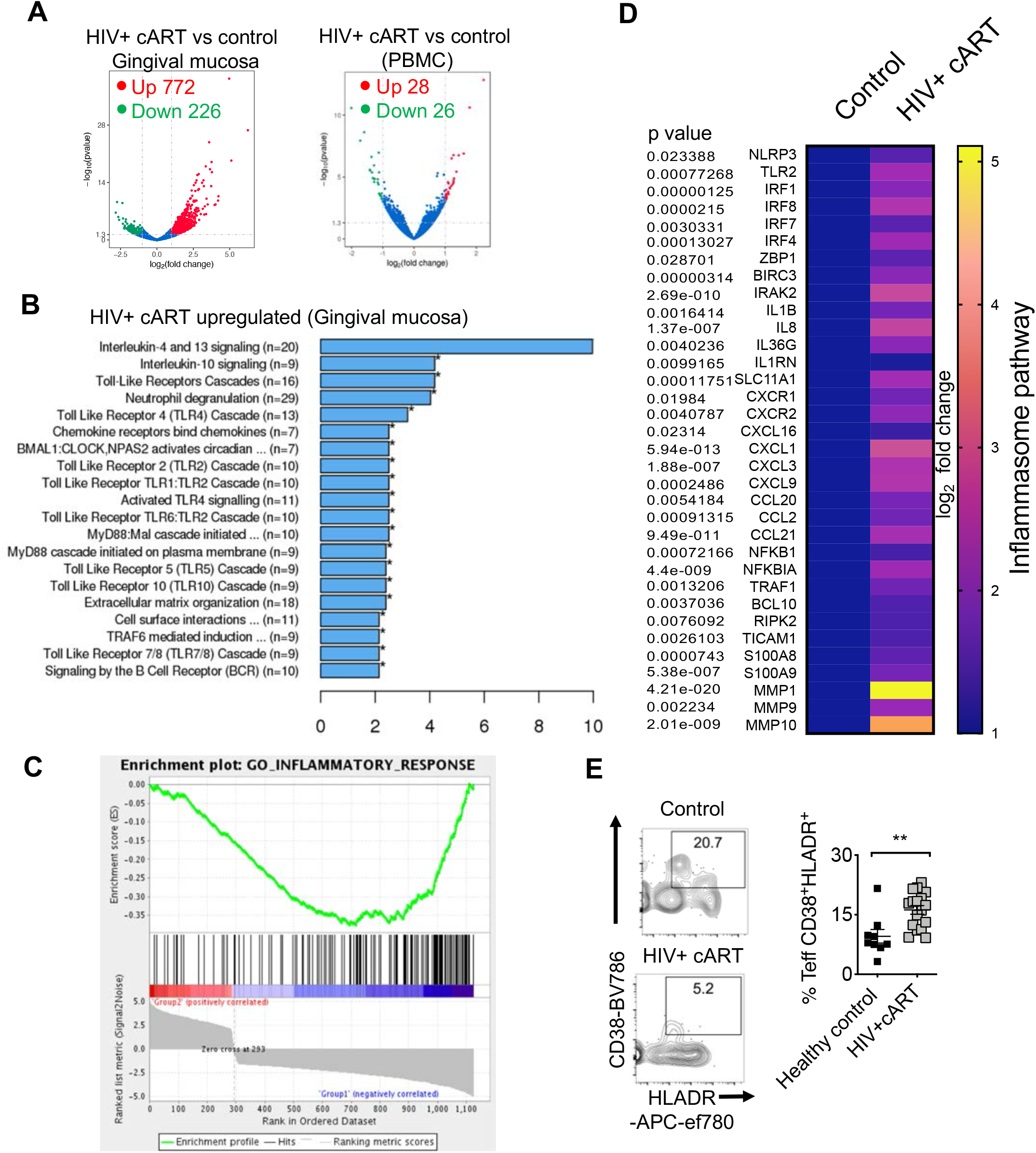
Transcriptomic profiling and flow cytometry analysis of oral mucosa in HIV+ patients. 46 HIV^+^ patients on cART treatment and 32 uninfected healthy controls were recruited (**Table 1**). RNA sequencing was performed in gingival tissues and PBMC collected from six randomly chosen age matched participants; Healthy uninfected control (n=3) and HIV+ cART (n=3), 2 males and 1 female in each group. Gingival cells were enriched for immune cells by reducing the epithelial cells through gradient centrifugation before transcriptome analyses. Volcano plots showing differential RNA expression in HIV+ cART *versus* healthy uninfected control groups in gingival mucosa (**A, left**) and PBMC (**A, right**). **B**) REACTOME pathway analysis of the genes upregulated in HIV+ cART gingival mucosa. **C**) Gene set enrich analysis (GSEA) was performed using the GSEA software (Broad Institute; http://www.broad.mit.edu/GSEA) employing the entire gene list generated from transcriptome analyses. This whole gene list was pre-ranked based on T-Score then uploaded to GSEA software. Inflammatory response signature genes were defined based on the gene sets in MSigDB. **D**) Heatmaps showing upregulation of inflammasome signature genes that were defined based on the published literature. Human oral intraepithelial and lamina propria leukocytes (HOIL) from gingival biopsies were processed for flow cytometry. **E**) Effector CD4 cells were gated as shown in **Fig.S1B,** and further on FOXP3 negative population. Contour plots (left) and statistics (right) showing the percentage of activated (CD38^+^ and HLADR^+^) effector CD4^+^ cells (n=20); (* P<0.05; Mann Whitney test).

### CD4^+^CD25^+^FOXP3^+^ cells are enriched in oral mucosa of HIV^+^ patients on therapy

Based on the upregulation of transcripts in the inflammasome pathway (**Fig.1D**), we then assessed IL-1β levels in HIV^+^ patients. We have previously shown increased IL-1β in lymphoid organs of HIV^+^ patients^37^, but the oral mucosa has not been examined. While IL-1β levels appeared to be lower (**Fig. S3A**), IL-6 levels were significantly higher in the saliva of HIV+cART patients (**Fig. S3B**). We also determined their expression in the supernatants of stimulated oral gingival immune cells *ex vivo*. These cells derived from HIV^+^ patients showed significantly elevated levels of secreted IL-1β and IL-6, corroborating with their inflammatory signature (**Fig.1C, D, 2A**). Given the role of microbial ligands in regulating mucosal cytokines, we hypothesized that dysbiotic oral microbiome may also be linked to alterations in cytokine levels and TLR signaling in oral mucosa of HIV^+^ patients ^38, 39^. Although D-dimers and lipoteichoic acid were not significantly different between the groups (data not shown), we found that salivary soluble TLR-2 proteins were significantly increased in HIV^+^ patients (**Fig.2B**). Interestingly, younger HIV^+^ patients (< 60) showed increased levels of sCD14 in their serum compared to young healthy controls(**Fig.S3C**). These features of inflammation, *i.e.* CD4 hyperactivation (CD38^+^HLA-DR^+^) and alterations in TLR-2 signaling led us to hypothesize that there might be a defect in immune regulation in oral mucosa of HIV^+^ patients^40, 41^. By first examining the transcriptome of gingival mucosa for the genes involved in promoting T_reg_ development and functions, we found that some of the T_reg_ transcripts were significantly enriched in oral mucosa of HIV^+^ patients (**Fig.2C**). Flow cytometry analyses of CD4, CD25, and FOXP3 expression also revealed that oral mucosal T_reg_ proportions were strikingly higher in the HIV^+^ group compared to the HIV-negative individuals (**Fig.2D, 2E, top**). However, there were no differences in T_reg_ percentages in PBMC between these two groups, showing that T_reg_ dysregulation was specific to the mucosa(**Fig.2D, 2E, bottom**). Because CD4^+^ T cells exhibited a hyperactivated phenotype (**Fig.1E**), we anticipated a lower frequency of FOXP3^+^ T cells in the HIV^+^ group but were surprised to find increased T_reg_ proportions in oral mucosa of HIV^+^ individuals. Increased TLR-2 signaling that we observed in HIV^+^ patients(**Fig.1B,1D, 2B**) can enhance FOXP3^+^ cell proliferation and alter the functions of T_reg_ and non-T_reg_ CD4^+^ T cells^42, 43^. It is known that oral complications such as periodontitis are of increased incidence and severity in HIV^+^ individuals even after suppressive HIV therapy. A majority of the HIV^+^ patients in our cohort had previous oral lesions. Therefore it is possible that generalized inflammation such as periodontitis contributes to T_reg_ dysregulation. To verify this possibility, we profiled FOXP3^+^ cells in gingival mucosa from both chronic and acute periodontitis non-HIV patients comparing them with healthy individuals. Although we found increases in Th17 cells in periodontitis non-HIV patients as shown previously^44^, there were no significant changes in the frequency of FOXP3^+^ cells in their gingiva (**Fig.S4**). These results show that previous inflammation does not by itself correlate or contribute to FOXP3^+^ T_reg_ cell enrichment in the oral mucosa. Taken together, these data raise the possibility that enrichment of FOXP3^+^ cells might be linked to the up-regulation of inflammasome and TLR/MyD88 signaling (**Fig. 1B, D**) and localized CD4^+^ T hyperactivation (**Fig.1E**) specific to the oral mucosa of HIV^+^ patients.

**Fig. 2:**
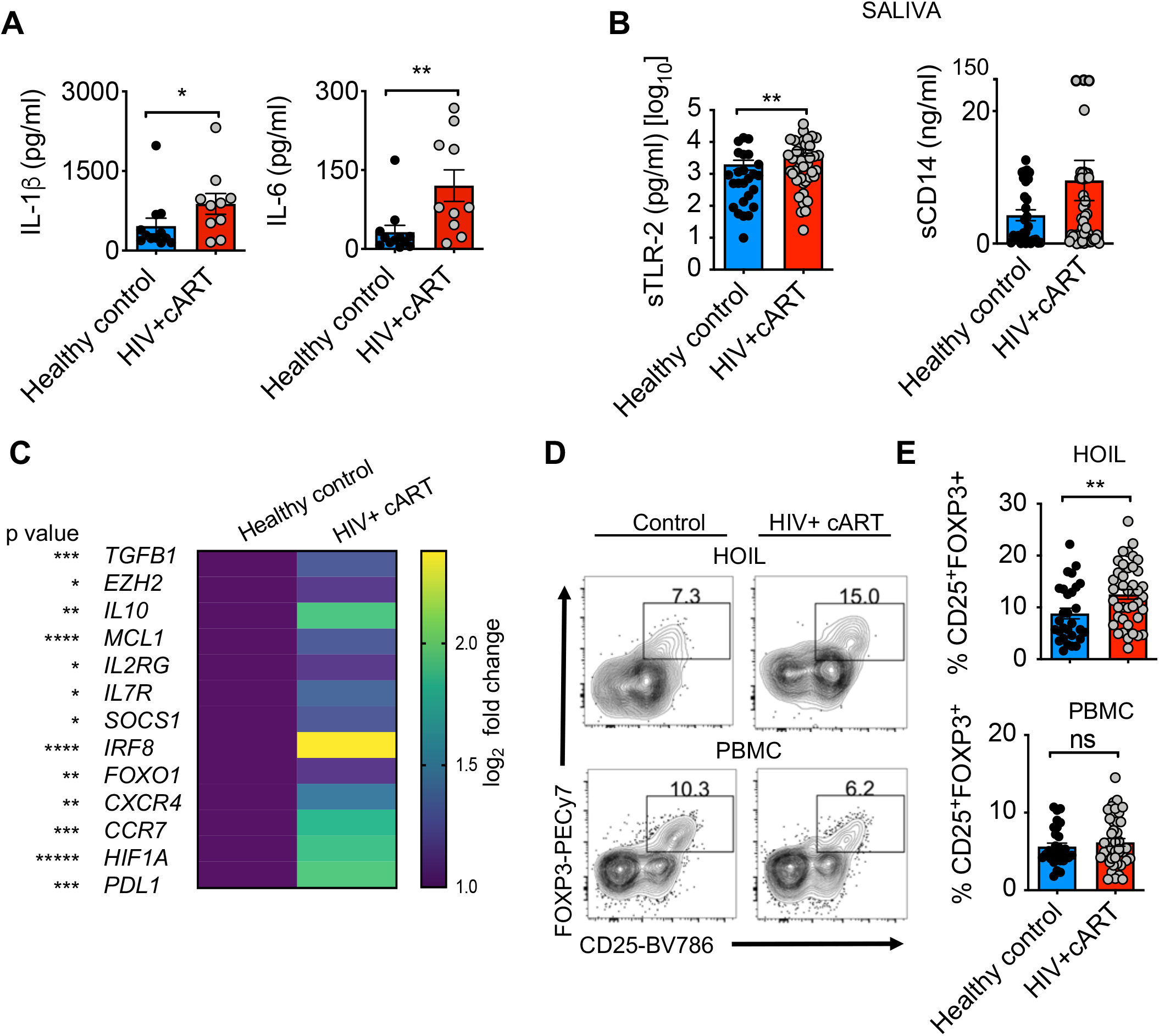
Inflammatory cytokines, sTLR-2 and CD4^+^CD25^+^FOXP3^+^cells are enriched in gingival mucosa of HIV^+^ patients on therapy. **A**) Cells from gingival mucosa were re-stimulated with PMA/Ionomycin for 4 hours and supernatants were collected for ELISA analyses of IL-1β (left) and IL-6 (right) (n=78). **B**) ELISA quantification of s-TLR-2 (left) and s-CD14 (right) levels in saliva. **C**) Transcriptome profiling was performed as in **Fig.1**. Heat maps of genes encoding literature curated Treg signature proteins differentially regulated in gingival mucosa. Flow cytometric analyses of CD45^+^CD3^+^CD4^+^ gated HOIL cells for CD25^+^FOXP3^+^ cell proportions, showing representative contour plots (**D**), and statistical analysis of Treg proportions (**E**) in HOIL (above) and PBMC (below). Mean values ± SEM are plotted. (* P<0.05; Mann Whitney test).

### HIV infection of oral MALT induces cell death and phenotypic changes in FOXP3^+^ cells

The oral mucosal system is composed of compartmentalized mucosa-associated lymphoid tissue (MALT) which includes palatine tonsils. The lymphoid environment of the tonsil oral MALT makes these tissues highly susceptible to infection and establishment of HIV reservoirs^45, 46^, however, CD4^+^ T cell dysfunction in relation to oral residual immune activation in cART treated patients has not been studied before. To obtain mechanistic details underlying T_reg_ alterations during HIV infection, we employed human tonsil cultures (HTC) derived from uninfected individuals. We hypothesized that this system would provide mechanistic insights into immune dysfunction in oral mucosa of HIV^+^ patients *in vivo.* First, we performed immunophenotyping of the tonsils that were obtained from tonsillectomy surgeries in children. As expected, examining the disaggregated tonsil cells in comparison with PBMC from independent healthy donors *ex vivo*, we found that tonsils harbored modestly lower frequencies of CD3^+^ T cells, NK and NKT cells, but significantly higher proportions of CD19^+^ B cells (**Fig.S5A, top, bottom, S5C**). Although tonsils had comparable levels of CD4^+^ T cells, they had reduced CD8^+^ T cells (**Fig.S5B, S5C**). As expected, 25-55% of CD4^+^ T cells were CXCR5^+^PD-1^high^ Follicular T helper cells (TFH) in tonsils (**Fig.S5D, top, S5E**). The overall proportions of CD4^+^FOXP3^+^cells were comparable to those in PBMC, but there were slightly lower proportions of CD25^+^FOXP3^+^ T follicular regulatory cells (TFR) and significantly higher proportions of CD45RO^+^ memory cells in tonsils when compared to PBMC (**Fig.S5D, middle, and bottom**). We stimulated whole tonsil cultures and infected them with the X4-tropic HIV-1 strain NLGNef ^47, 48^. We characterized the productively infected cells by examining the GFP expressing cells (**Fig.S6A**). As shown before ^46^, 95% of the productively infected cells were of germinal center TFH phenotype (**Fig.S6B, top**), but co-expressed FOXP3 and CD45RO(**Fig.S6B, middle**). Though T_regs_ have been previously shown to be an HIV-permissible population^49^, our results show that these p24 expressing cells also co-express CD45RO (**Fig.S6B, middle, bottom**). They also expressed higher levels of CXCR4, with many co-expressing CCR5, consistent with a previous report on tonsillar CD4^+^ T cells (^50^, data not shown). Examining the FOXP3^+^ cell fraction more closely, PD-1^high^CXCR5^+^CD45RO^+^ cells, which were consistent with the memory phenotype germinal center follicular regulatory cells (TFR), displayed a significant high-permissibility to HIV infection(**Fig.S6C, D**). As shown previously^45^, these TFR showed higher permissibility to both X4- and R5 tropic viruses (data not shown). The expression of surface and intracellular IL-10R in jejunum lamina propria, but not jejunum intraepithelial T cells during late acute SIV infection^51^. Therefore we examined the expression of the IL-10R receptor but did not find changes in IL-10R expression with and without anti-retroviral inhibitor in HIV-infected CD4^+^ T cells (**Fig.S6E**).To distinguish whether these permissible FOXP3^+^ cells were pre-existing tonsil T_regs_ or induced during TCR stimulation, we purified CD4^+^ cells and CD4+CD25^+^CD127^low^ cells and infected them with HIV. We found that FOXP3^+^ cells were highly permissible in both of the cultures, although purified T_regs_ harbored a significantly higher frequency of GFP^+^ cells (**Fig.3A**). Similar results were obtained even in the absence of TCR activation, as tonsil cells do not require exogenous stimulation prior to infection (data not shown; ^45^). These results confirm previous studies showing high permissibility of FOXP3^+^ cells to HIV infection^45, 49^. Moreover, we found that the proportions of FOXP3^+^, as well as FOXP3^negative^ IL-17A^+^ and IFN-γ^+^ effector CD4^+^ populations, are decreased in HIV infected tonsils (**Fig.3B, S7 A, B**), showing that CD4^+^ cells are also highly susceptible to cell death during acute HIV infection (**Fig.3C**). This is consistent with previous results on HIV-mediated apoptotic and pyroptotic CD4^+^ T cell depletion^50, 52^. Interestingly, we found that the frequency of PD-1^hi^IFN-γ^+^ cells among the viable FOXP3^+^ population consistently increased with HIV^+^ infection (**Fig.3D; S8;** staining controls). Although this population was productively infected (**Fig.S9A**), it expressed high levels of BCL-2 and was more resistant to cell death compared to PD-1^low^ cells (**Fig.3E, top**). Further characterization of this population revealed that they expressed high levels of CD25(**Fig.S9B**), IL-1 family receptors such as IL-1R, IL-33R (Suppression of Tumorigenicity 2; ST-2), and amphiregulin (AREG) (**Fig.3E, middle and bottom**), resembling activated tissue T_regs_. While this population had slightly lower levels of BCL-6, they expressed IL-10 (**Fig.3E**) and B lymphocyte-induced maturation protein 1 (BLIMP1), characteristic of effector TFR cells in germinal centers and tissue T_regs_ (**Fig.S9C**)^53^. Taken together, these data revealed that although HIV infection led to the loss of CD4^+^ T cells, it resulted in an increase of a unique population of PD-1^hi^IFN-γ^+^ AREG^+^FOXP3^+^cells that survived the infection and might contribute to immune dysfunction.

**Fig. 3.**
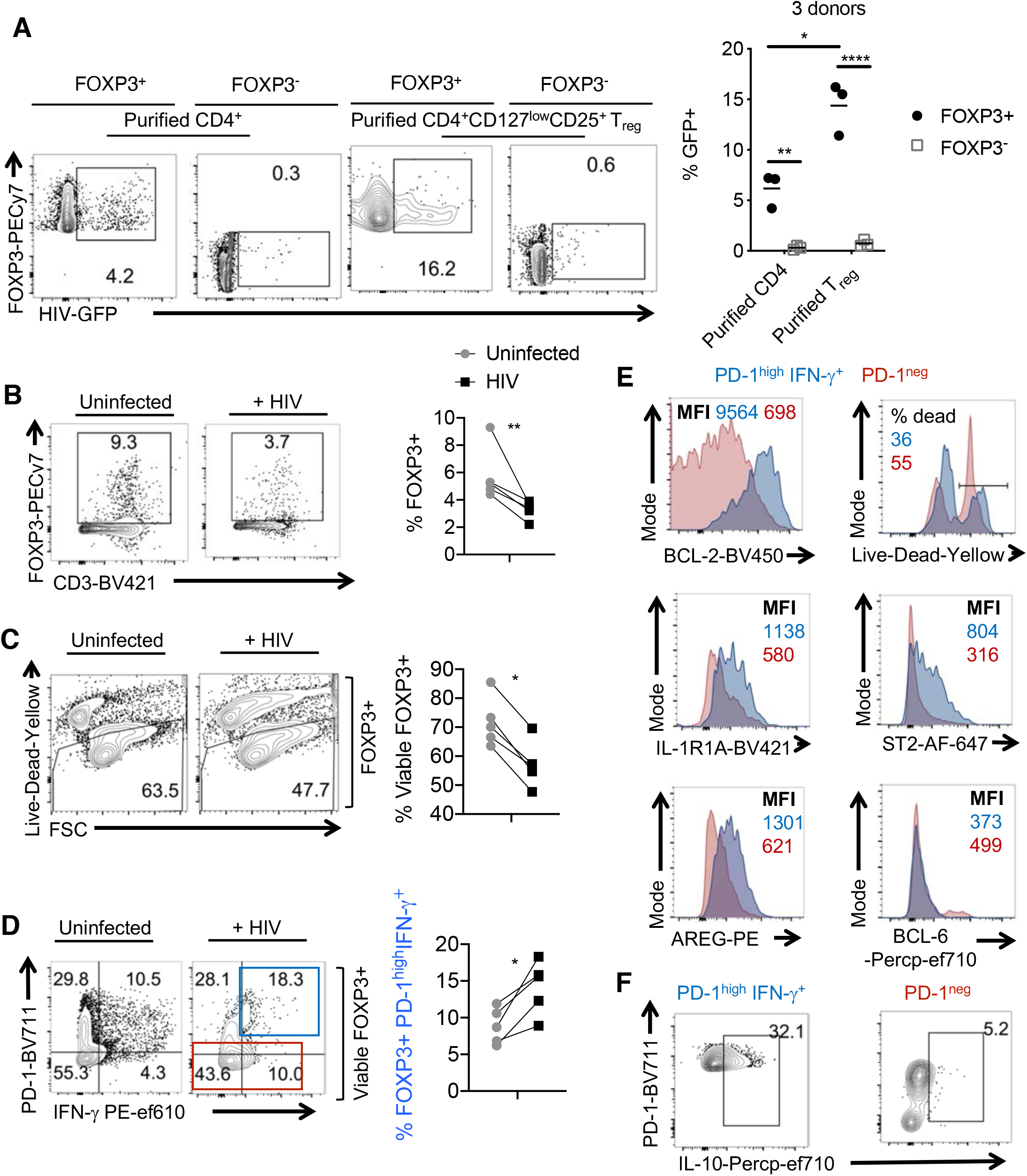
HIV infection reduces FOXP3^+^ cells but increases the proportions of PD-1^high^IFN-γ^+^ cells among FOXP3^+^ *in vitro*. **A**) Purified tonsil CD4^+^ T cells (∼91% purity) or CD4^+^ CD25^+^ CD127^low^ Treg cells (> 88% FOXP3^+^) were TCR activated and infected with HIV as described in methods. GFP was assessed in FOXP3^+^ (left) or FOXP3^-^ (right; gated on Foxp3^neg^ cells) fractions 48 hours post-infection. Representative flow cytometric data (left) and statistical analyses from 3 independent tonsil donors (right) are shown. **B-F**) TCR activated whole human tonsil cultures (HTC) were infected with HIV and allowed to expand with IL-2 for 6 days. Flow cytometric analyses of CD3^+^FOXP3^+^ cells pre-gated on CD8 negative cells (**B**), viability of CD3^+^ CD8 negative FOXP3^+^cells (**C**), PD-1 and IFN-γ expression in viable CD3^+^FOXP3^+^CD8 negative cells (**D**), with respective statistical analyses from 5 experiments (right) are shown. Mean values ± SEM are plotted. **E,F**) Flow cytometric plots showing the expression of indicated proteins in PD-1^high^ and PD-1^low^ populations gated in (**D**) in HIV infected HTC. At least 5 independent experiments showed similar results.

### Blocking subsequent rounds of infection and cell death increased the proliferation of PD-1^hi^IFN-γ^+^FOXP3^+^cells

We further characterized the conditions under which these FOXP3^+^cells were induced during HIV infection and tested whether mechanisms underlying HIV-induced cell death might play a role. Therefore, we aimed to block cell death by inhibiting HIV replication and caspase activation after the onset of initial cell death. To this end, we added reverse transcriptase inhibitor Efavirenz and pan-caspase inhibitor 28 hours after HIV infection. We found that both were able to increase the overall viability of CD4^+^T cells including FOXP3^+^ cells (**Fig.4A, B,** data not shown). Interestingly, while the proportions of PD-1^hi^IFN-γ^+^ FOXP3^+^ cells were partially reduced by these inhibitors, their absolute cell numbers significantly increased in the cultures (**Fig.4C**). These data show that PD-1^hi^IFN-γ^+^ FOXP3^+^ cells that were induced during initial HIV infection had a survival advantage and likely expanded in the presence of these inhibitors in oral MALT. Consistent with this notion, while PD-1^low^FOXP3^+^ cells did not proliferate much, the percentage and absolute numbers of Ki-67^+^ cells were higher in PD-1^hi^FOXP3^+^ cells in the presence of these inhibitors (**Fig 4D**). Collectively, these data highlight that initial HIV infection is sufficient for PD-1^hi^IFN-γ^+^ FOXP3^+^ cell accumulation, and these cells are not abolished with the antiviral drug treatment. Instead, blocking HIV replication and HIV induced cell death after the initial rounds of HIV infection promoted the proliferation of PD-1^hi^IFN-γ^+^ FOXP3^+^ cells that were rescued from cell death.

**Fig. 4.**
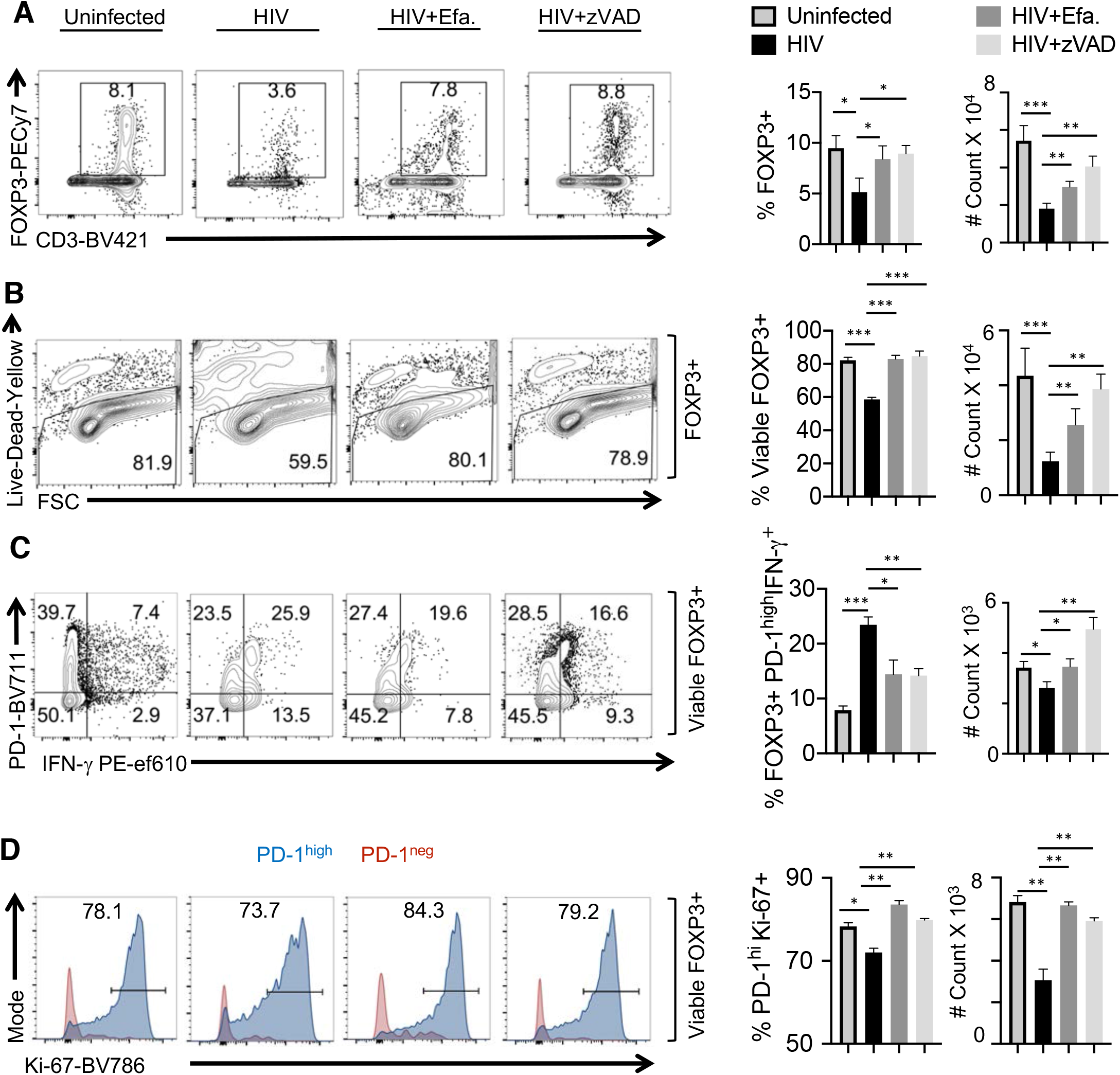
Blocking subsequent rounds of infection and cell death increased the proliferation of PD-1^hi^IFN-γ^+^FOXP3^+^cells. Whole HTC was activated with TCR stimulation, infected with HIV and allowed to expand in the presence of TGF-β1 (10 ng/ml) and IL-2 (100 U/ml) for 6 days. Viral inhibitor Efavirenz (50 nM) or cell death/ pan-caspase inhibitor z-VAD (10 μM) was added 28 hours post-infection as described in methods. Flow cytometry acquisition was done with constant time for all the samples. Percentage of CD3^+^FOXP3^+^ cells pre-gated on CD8 negative cells (**A**), Viability of FOXP3^+^ cells pre-gated on CD3^+^CD8 negative cells (**B**), PD-1 and IFN-γ expression in viable CD3^+^ CD8 negative FOXP3^+^ cells(**C**), Ki-67 expression in viable PD-1^high^ and PD-1^low^ FOXP3^+^ populations (**D**) are shown. Representative contour plots (left), statistical analyses of proportions of the cells (middle) and statistical analyses of the absolute cell counts (right) are shown (2way ANOVA, multiple comparison; alpha= 0.05*).

### PD-1^hi^IFN-γ^+^ FOXP3^+^ cell accumulation is associated with the expression of IL-1β-dependent AKT1 signaling and enhanced by TLR-2 ligands in the context of HIV infection

We and others have previously shown that direct and indirect TLR-2 signaling in FOXP3^+^ cells can induce proliferation impacting their functions^40^. Moreover, results from HIV^+^ patients that showed TLR-2 pathway upregulation and cytokine inflammatory pathways in the oral mucosa (**Fig.1, 2**) led us to hypothesize that TLR-2 signaling is involved in PD-1^hi^IFN-γ^+^ FOXP3^+^ cell induction. There is copious evidence that HIV^+^ patients have episodes of recurring oral *Candida* infections and periodontitis despite therapy (**Table 1**), which might contribute to the enrichment of transcripts involved in TLR signaling in their oral mucosa (**Fig.1, 2**)^15, 54, 55^. To this end, we determined the effect of lipopolysaccharide (LPS) and TLR-2 ligands such as *Candida* (heat-killed germ tube (HKGT) and *Porphyromonas gingivalis* (PG-LPS) on purified tonsil CD4^+^ cells in the context of HIV infection. While HKGT moderately increased PD-1^hi^IFN-γ^+^ FOXP3^+^ cells, these ligands did not alter cell viability or expansion of PD-1^hi^IFN-γ^+^ FOXP3^+^ cells in uninfected cultures (**Fig.5A**). However, in HIV-infected cultures, these ligands promoted a significant increase in PD-1^hi^IFN-γ^+^ FOXP3^+^ cells, as well as AREG expression in FOXP3^+^ cells (**Fig.5A, left and right, S10**). We saw consistent results even in the absence of TCR stimulation of CD4^+^ T cells(**Fig. S11A, B**). To determine the mechanism underlying the accumulation of PD-1^hi^IFN-γ^+^FOXP3^+^ cells and AREG expression in these cells, we examined the cytokine production in cultures. A previous study has shown that HIV induces the secretion of pyroptosis-related cytokine IL-1β in CD4^+^ T cells^50^. Based on the upregulation of IL-1β and IL-6 in oral mucosa of HIV^+^ patients (**Fig.2A**) and the role of IL-1 family cytokines in promoting AREG expression^56, 57^, we examined the effect of IL-1β, IL-33, and IL-6 in HIV infected tonsil CD4^+^ T cell cultures. ELISA quantification demonstrated that HIV infection elevated the levels of mature IL-1β and AREG, which were further increased when CD4^+^ T cells were stimulated with TLR-2 ligands (**Fig. 5B**). While HIV infection did not alter IL-33 and IL-6, it enhanced IL-1β, which was further upregulated by TLR-2 ligands (**Fig. S11C, D**). Induction of mature IL-1β is likely caspase-1 dependent, and this cytokine can function in CD4 intrinsic and phosphatidylinositol-3-OH kinase (PI-3K)/AKT1 dependent manner in effector CD4^+^ T cells^50, 58–61^. Moreover, AKT-1 activation/phosphorylation and FOXO3 repression promote activated T_reg_ cell accumulation in tissues^62^. Indeed, we found that HIV infection was able to activate caspase-1, as measured by its phosphorylation in FOXP3^+^ cells (**Fig. S12**). TLR-2 ligands further enhanced caspase-1 activation almost to the levels of Nigericin, an IL-1/inflammasome, and pyroptosis activator (**Fig. S12**).

**Fig. 5.**
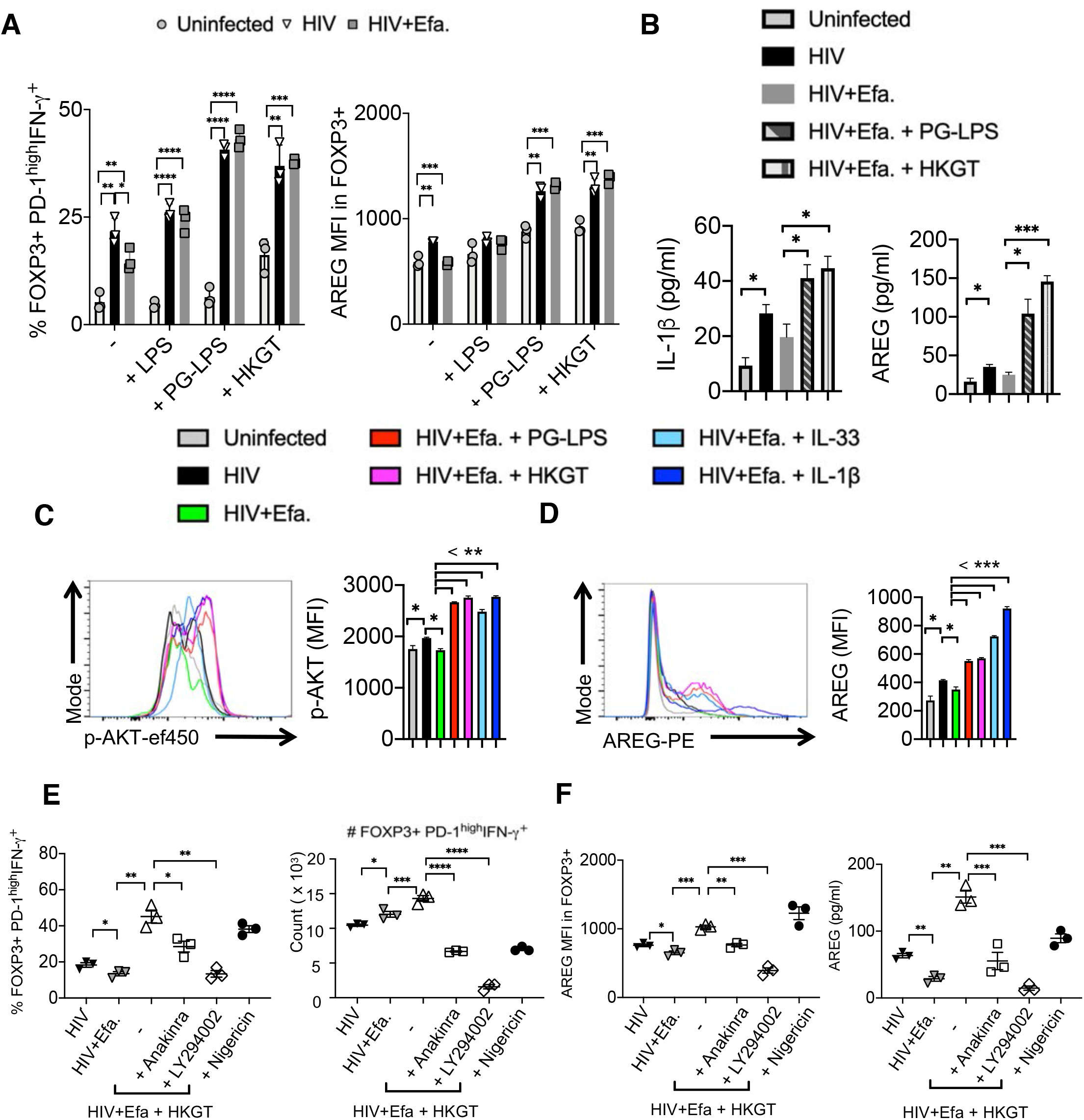
PD-1hiIFN-γ+ FOXP3^+^ cell induction is associated with expression of IL-1β-dependent AKT1 signaling and enhanced by TLR-2 ligands in the context of HIV infection. Purified tonsil CD4^+^ T cells (∼93% purity) were TCR activated, infected with HIV and allowed to expand in the presence of TGF-β1 (10 ng/ml) and IL-2 (100 U/ml) for 6 days post-infection unless otherwise noted. Efavirenz (50 nM), LPS(10μg/ml), PG-LPS (5 μg/ml), HKGT ( 10^6^/ml), IL-1β (20 ng/ml), IL-33 (20 ng/ml), Anakinra (10 μg/ml), LY294002 (10 μM) and Nigericin (10nM) were added as indicated, 36 hours post infection. **A**) PD-1 and IFN-γ (left) and AREG (right) expression in FOXP3^+^cells. **B**) ELISA quantification of IL-1β (left) and AREG (right) in cell culture supernatants collected on day 3 post infection. p-Akt (**C**) and AREG (**D**) expression in FOXP3^+^ cells. (**E**) Percentage and absolute cell numbers of PD-1^hi^IFN-γ^+^ FOXP3^+^ cells in CD4^+^ population. (**F**) AREG expression in FOXP3^+^ cells (left) and ELISA quantification of AREG (right), 6 days post infection. Data are representative of at least 3 independent experiments.

Next, we examined whether TLR-2 ligands or IL-1β can promote PD-1^hi^IFN-γ^+^FOXP3^+^ cells and AREG expression induction from naïve T_regs_ in the context of HIV infection. About 82-92% of FOXP3^+^cells in tonsils are of CD45RO^+^CD62L^low^ phenotype(**Fig. S13A**). As expected, CD45RO^neg^ naïve T_regs_ were CD62L^high^ and PD-1^neg^(**Fig. S13A**). To address whether PD-1^high^T_regs_ can be induced from naïve T_regs_, we isolated CD45RO^neg^ naïve T_reg_ cells from tonsils and infected them with HIV infection in the presence of TLR-2 ligand. The frequency of PD-1^high^ cells and AREG expression are much lower in these cultures when compared to non-purified CD4 T cell cultures (compare **Fig. S13B** with **Fig.5A**). Also, pre-existing CD45RO^+^T_reg_ cells have increased BCL-2 expression compared to CD45RO^neg^ FOXP3^+^ cells (**Fig. S13C**). Additionally, we also sorted PD-1^+^T_reg_ and PD-1^neg^ T_reg_ cells from tonsils and examined the expression of secondary markers such as IFN-γ and AREG with and without infection (**Fig. S14**). Consistent with our hypothesis and the results in Fig.3, purified PD-1^+^T_reg_ cells showed higher IFN-γ and AREG expression, compared to PD-1^neg^ T_reg_ cells (**Fig. S14A, B**). They also showed higher expression of Ki-67, HIV-GFP, and BCL-2 expression than PD-1^neg^ T_reg_ cells(**Fig.S14C, D, E**). These data support the notion that PD-1^+^T_regs_ although has high infection susceptibility, may intrinsically survive and proliferate better with HIV infection, leading to the accumulation of dysfunctional T_regs_. Interestingly, a small proportion PD-1^neg^T_reg_ population can also upregulate PD-1 and IFN-γ in the context of HIV infection (but not TLR-2 stimulation alone). These induced cells also show higher proliferation with TLR-2 and IL-1β stimulation in the context of HIV infection, but not as much as purified PD-1^+^T_regs_(**Fig. S14C**). However, PD-1^hi^IFN-γ^+^AREG^high^FOXP3^+^ cells could not be induced from naïve CD4^+^ T cells during HIV infection (data not shown). Taken together, these data show that while pre-existing PD-1^+^FOXP3^+^ cells might contribute more to the accumulation of dysfunctional T_regs_, naïve PD-1^neg^FOXP3^+^ cells can also be induced to become PD-1^high^ cells expressing high levels of IFN-γ and AREG.

Finally, we determined the ability of IL-1 cytokines and TLR-2 ligands to activate AKT kinase downstream in the PI-3K pathway. Because of the ability of IL-1 cytokines to upregulate AREG in tissue T_regs_ ^56, 57, 63^, we also examined AREG expression in FOXP3^+^ cells. While HIV was able to increase the accumulation of PD-1^hi^IFN-γ^+^ FOXP3^+^ cells and moderately induce phosphorylation of AKT and AREG expression, TLR-2 ligands, IL-1β, and IL-33 significantly enhanced AKT phosphorylation and AREG expression in FOXP3^+^ cells (**Fig.5C, D, S15**). However, there were neither alterations in STAT-3 phosphorylation nor an effect of IL-6 on inducing PD-1^hi^IFN-γ^+^ FOXP3^+^ cells (data not shown). Based on these observations, we next investigated the function of IL-1β–induced AKT1 signaling pathway in promoting PD-1^hi^IFN-γ^+^ FOXP3^+^ cells and AREG expression in FOXP3^+^ cells. Both drugs, the inhibitors of IL-1β signaling (Anakinra) and PI-3K/ AKT1 (LY294002), significantly reduced the percentage and absolute cell numbers of HIV-induced PD-1^hi^IFN-γ^+^ FOXP3^+^ cells in tonsil cultures(**Fig.5E, S16**). Also, IL-1β and AKT1 inhibition downmodulated AREG expression in FOXP3^+^ cells (**Fig.5F**), suggesting synergistic roles of HIV, TLR-2 ligands, and the IL-1β in altering FOXP3^+^ cells in an AKT1 dependent fashion during HIV infection.

### PD-1 signaling stabilizes the expression of FOXP3 and AREG by downmodulating asparaginyl endopeptidase (AEP)

In the above experiments, we observed that IL-1β was able to promote PD-1 expression in FOXP3^+^ cells in a manner dependent on AKT1 activation (**Fig.S14, S16; y-axis**). This led us to interrogate whether PD-1 signaling directly regulated HIV-induced PD-1^hi^IFN-γ^+^ FOXP3^+^ cells. PD-1 has been previously shown to modulate AEP, an endo-lysosomal protease implicated in antigen processing and FOXP3 expression^64, 65^. Therefore, we further characterized the PD-1^high^ and PD-1^low^ FOXP3^+^ cells in HIV-infected CD4^+^ T cell cultures in the presence of Efavirenz added 28 hrs after infection. Although PD-1^high^ FOXP3^+^ cells had slightly higher expression of AEP, levels of phosphorylated AEP (pAEP), the active form of AEP enzyme, were precipitously lower than in PD-1^low^ FOXP3^+^ cells (**Fig.6A, 1^st^ 2 panels**). Also, these PD-1^high^FOXP3^+^ cells had higher expression (higher MFI) of FOXP3 compared to PD-1^low^FOXP3^+^ cells (**Fig.6A, 3^rd^ panel**). Concurrent with their enhanced survival and proliferation, PD-1^high^ FOXP3^+^ cells had elevated expression of BCL-2 and Ki-67 (**Fig.6B**). Engaging PD-1 using PD-1 ligand-Fc (PDL-1-Fc), or inhibiting AEP using an inhibitor increased FOXP3 expression in HIV infected CD4^+^ cells(**Fig.6C**), suggesting that active PD-1 signaling in the context of IL-1β is involved in the stability of FOXP3 expression during HIV infection. PD-L1-Fc and AEP inhibition heightened the frequency and absolute numbers of PD-1^hi^IFN-γ^+^ FOXP3^+^ cells(**Fig.6D**) showing that the PD-1-AEP axis is critical for the survival and proliferation of PD-1^hi^FOXP3^+^ cells. Moreover, PD-1 engagement and AEP inhibition promoted the expression of AREG in PD-1^+^FOXP3^+^ cells (**Fig.6E**). Purified PD-1^neg^ cells that were activated and infected as in **Fig.S14**, lose FOXP3, which further confirms that PD-1 is required for Foxp3 retention (**Fig.S17**). Altogether, these results showed that direct PD-1 signaling enhances FOXP3 and AREG expression by inhibiting AEP in the context of IL-1β expression during HIV infection.

**Fig. 6.**
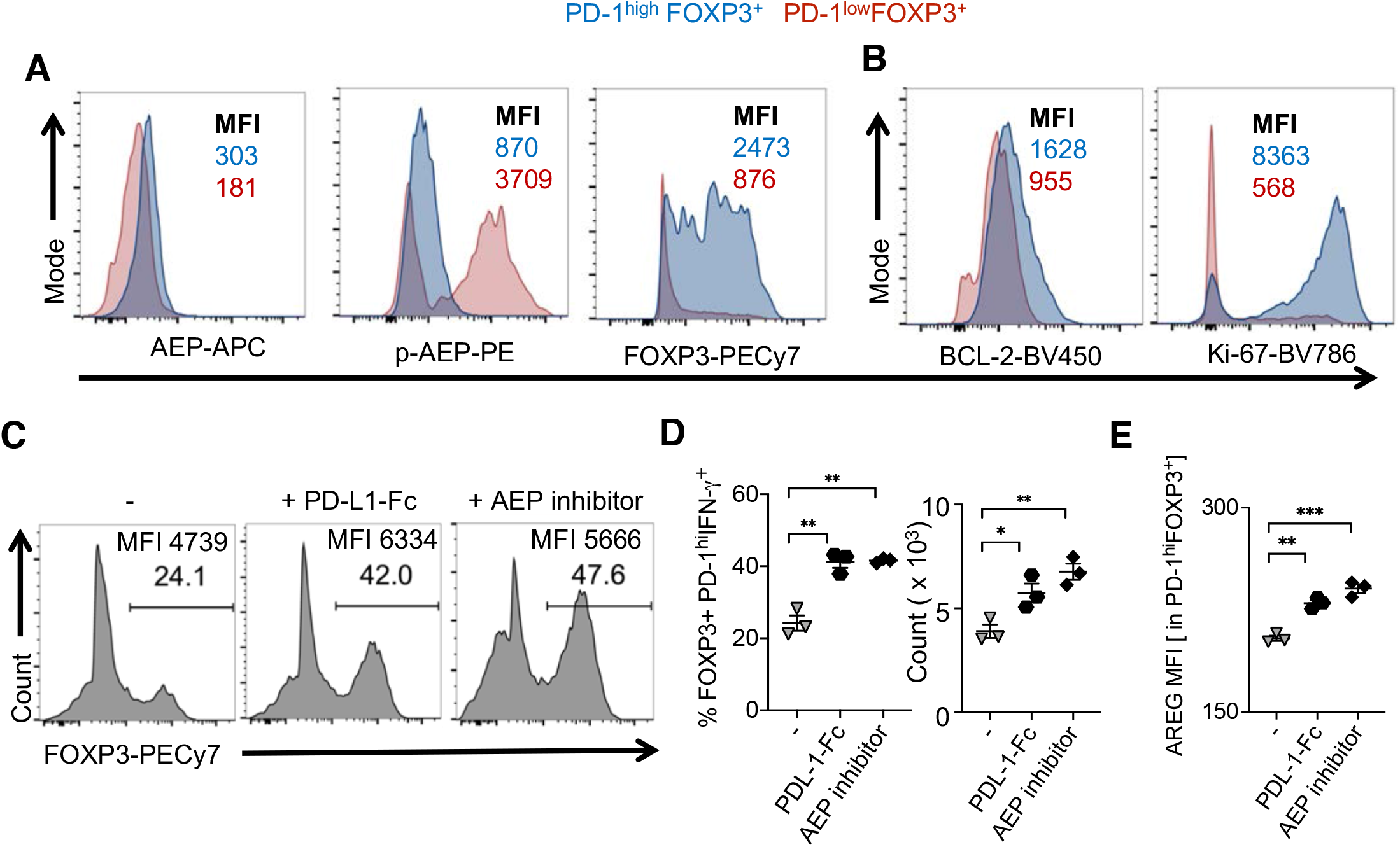
PD-1 ligation downmodulates asparaginyl endopeptidase (AEP) and stabilizes the expression of FOXP3 and AREG. CD4+ T cells were stimulated as in **Fig.5**. **A,B**) AEP, pAEP, FOXP3,BCL-2 and Ki-67 staining in PD-1^high^FOXP3^+^ (blue) and PD-1^low^FOXP3^+^ cells 6 days post-infection. Some CD4^+^ T cells stimulated and infected as above were moved to a plate coated with recombinant human PD-L1/B7-H1 Fc chimera (5 μg/ml), or treated with AEP inhibitor (10 μM) 36 hours after infection. Percentage of FOXP3^+^cells in CD4^+^ population and FOXP3 MFI on FOXP3^+^ gated cells (**C**), Percentage and absolute cell numbers of PD-1^hi^IFN-γ^+^ cells in FOXP3^+^ population (**D**), and AREG expression in FOXP3^+^ cells (**E**), as determined by flow cytometry analyses. Results represent triplicate experiments with similar results.

### PD-1^high^FOXP3^+^IFN-γ^+^ cells from HIV infected cultures have little or no suppressive activity

Next, we explored the function of PD-1^hi^IFN-γ^+^FOXP3^+^AREG^high^ cells that were induced during HIV infection *in vitro* and compared them with purified naïve CD4^+^CD25^+^CD127^low^ FOXP3^+^ cells activated and infected in a similar manner. To this end, we activated and infected tonsillar CD4^+^ T cells or purified CD4^+^CD25^+^CD127^low^ FOXP3^+^ T_regs_ as before (**Fig.7A, top**) and analyzed the proportion of PD-1^hi^IFN-γ^+^ within the FOXP3^+^ population. Interestingly, purified T_regs_ harbored significantly lower proportions of PD-1^hi^IFN-γ^+^cells (**Fig.7A, bottom, 7B**), suggesting that PD-1^hi^IFN-γ^+^FOXP3^+^AREG^high^ cells are derived preferentially from conventional CD4^+^T cells and T_regs_ that upregulate and maintain FOXP3 during activation. Next, we purified the PD-1^hi^CD25^+^ cells from CD4^+^ cell cultures which were HIV infected in the presence of Efavirenz and examined their suppressive activity. Cells purified from these cultures were ∼76-88% FOXP3^+^ and >50% IFN-γ^+^ positive (**Fig.S18**). We evaluated their ability to suppress the proliferation of CD4^+^ T cells by co-culturing them with cell-trace labeled activated tonsil CD4^+^CD25^neg^ responder T cells (T_resp_) from the same donor, as shown previously^58^. As controls, we had CD4^+^CD25^neg^ activated alone, and in co-cultures with purified T_regs_ that were activated and infected with HIV in the presence of Efavirenz. As expected, purified T_regs_ reduced the frequency of proliferating T_resp_ cells. However, at all time-points after activation, PD-1^hi^CD25^hi^ FOXP3^+^ cells did not affect the proliferation of T_resp_ cells in the co-cultures (**Fig.7C, 7D**), showing that PD-1^hi^CD25^hi^ FOXP3^+^ cells induced during HIV infection were dysfunctional in the context of their direct suppression of CD4^+^ T cell survival or proliferation. Also, blocking IFN-γ using an α-IFN-γ antibody (10 μg/ml) in PD-1^high^ T_reg_ co-cultures did not restore their suppressive capacity *in vitro* (data not shown). Collectively, these data show that PD-1^hi^IFN-γ^+^FOXP3^+^AREG^high^ cells derived from HIV-infected cultures do not suppress CD4^+^ T cells *in vitro*.

**Fig. 7:**
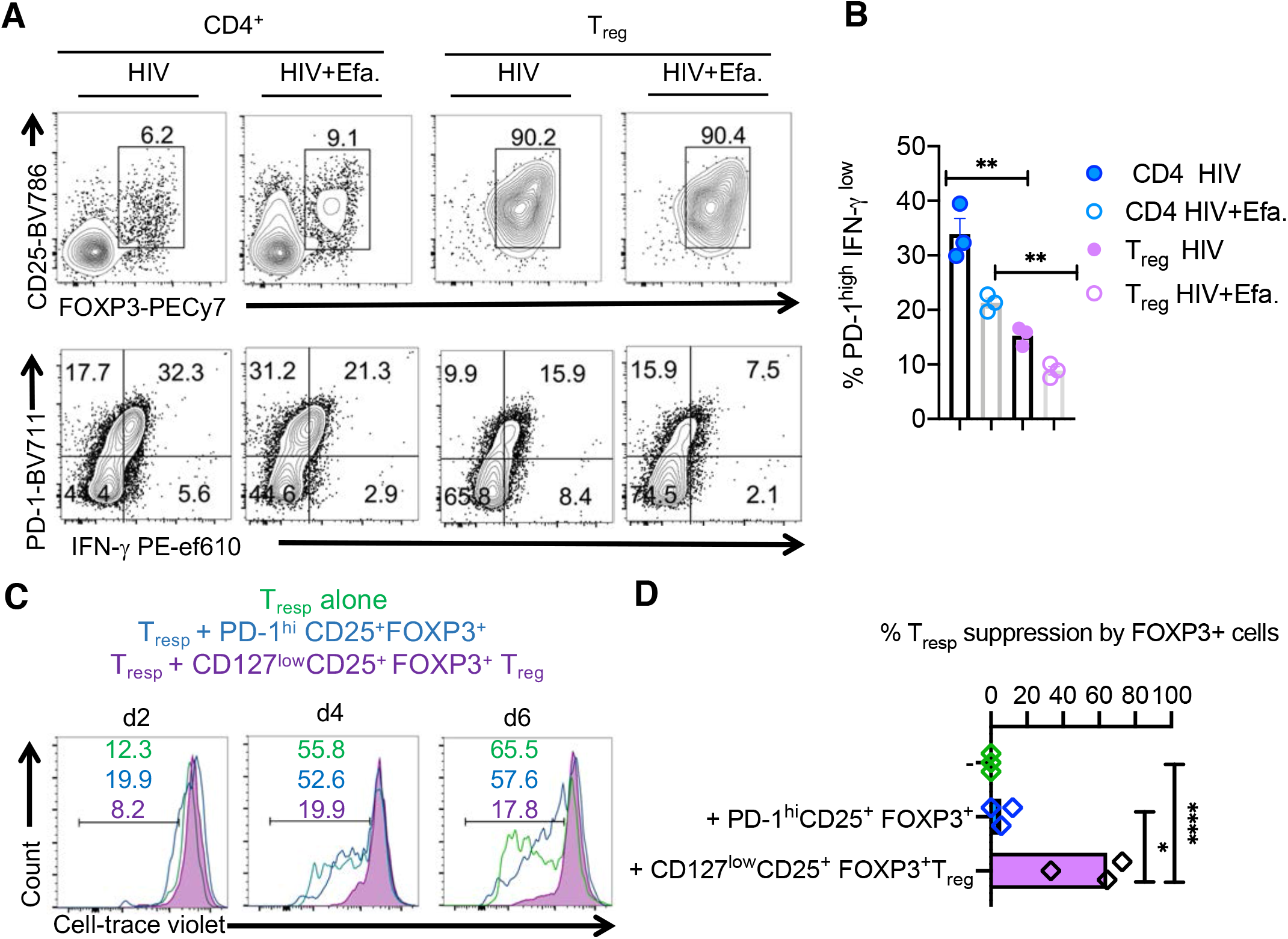
PD-1^+^ FOXP3^+^ cells from HIV infected cultures have little or no suppressive activity. Purified CD4^+^ T cells and Tregs were stimulated and infected as in methods. (**A**) CD25 and FOXP3 expression in all cells in the cultures (above) and PD-1 and IFN-γ expression in CD25^+^FOXP3^+^ cells (below) at 96 hours post infection (**B**) Statistical analyses of PD-1^hi^IFN-γ^+^ cells in FOXP3^+^ population from these two cultures. (**C**) PD-1^hi^CD25^+^ cells were purified from HIV-infected CD4 cultures using sequential sorting of PD-1-PE^+^ cells and CD25^high^ Treg cells using STEMCELL technology PE isolation and CD25^+^Treg isolation kits, and were used in co-cultures with cell-trace violet labelled responder T cells (Tresp) at ratio 1:1. As controls, Tresp cells were cultured alone or co-cultured with purified naïve CD127^low^CD25^+^Tregs that were sequentially sorted to remove CD45RO^+^CD4^+^cells using human CD45RO kit (Miltenyi) and purify CD25^high^Treg cells using STEMCELL technology CD25^+^Treg isolation kits. These control Tregs were also previously stimulated and infected the same manner (purple) before co-culture with Tresp. Tresp proliferation was determined by cell-trace dye dilution in PD-1^hi^CD25^+^co-culture (blue), control Treg co-cultures (purple), or those cultured alone (green) (**D**) Statistical analyses of % Tresp suppression mean values from three independent experiments showing similar results (* P<0.05; Mann Whitney test).

### The abundance of PD-1^hi^CD25^hi^ IFN-γ^+^AREG^hi^ FOXP3^+^ cells correlates with oral mucosal CD4 hyperactivation in oral mucosa of HIV+ patients

Although oral mucosa of HIV^+^ patients had a significantly higher frequency of FOXP3^+^ cells (**Fig. 2D, E, 8A**), because of the associated inflammatory signature and elevated IL-1β signaling (**Fig. 1D-F, 2A**) we hypothesized that FOXP3^+^ cells accumulating in oral mucosa of HIV^+^ patients may also be dysfunctional. To test this notion, we evaluated the expression of dysfunctional markers, PD-1 and IFN-γ. Although only a small proportion (∼ 6-9%) of CD4^+^CD25^+^FOXP3^+^ cells from healthy controls expressed PD-1, about 14-19% of them expressed PD-1 in HIV^+^ patients (**Fig. 8B, y-axis, E, F**). Concurring with the results from oral MALT HIV infection *in vitro* (**Fig.3**), HIV^+^ patients on cART also had a significantly higher percentage of PD-1^high^FOXP3^+^cells, co-expressing IFN-γ, IL-10, and AREG in the oral mucosa (**Fig.8B, 8C, upper right quadrants, 8D, E, G, H; S19; FMO controls**). AREG levels in saliva were also found to be elevated in HIV^+^ patients on cART when compared to healthy control individuals (**Fig.8I**). Although the FOXP3^+^ cells fit the profile of dysfunctional FOXP3^+^ cells incapable of CD4+ T cell suppression (**Fig.7**), we could not directly evaluate the suppressive function of patient oral mucosal FOXP3^+^ cells because of technical limitations. Nonetheless, the frequencies of PD-1^+^IFN-γ^+^FOXP3^+^ cells and salivary AREG levels showed a significant positive correlation with CD4^+^ hyperactivation (CD38 and HLA-DR co-expression) in the oral mucosa (**Fig.8J, K**), suggesting that FOXP3^+^ cells might indeed be impaired in their ability to suppress CD4^+^ T cells. Taken together, these data from oral gingival mucosal cells of HIV^+^ patients on cART substantiate the results from *in vitro* tonsil experiments and demonstrate that dysfunctional PD-1^+^AREG^+^ FOXP3^+^ cells strongly correlate with CD4 hyperactivation and contribute to the dysregulated immune landscape in treated HIV^+^ patients.

**Fig. 8.**
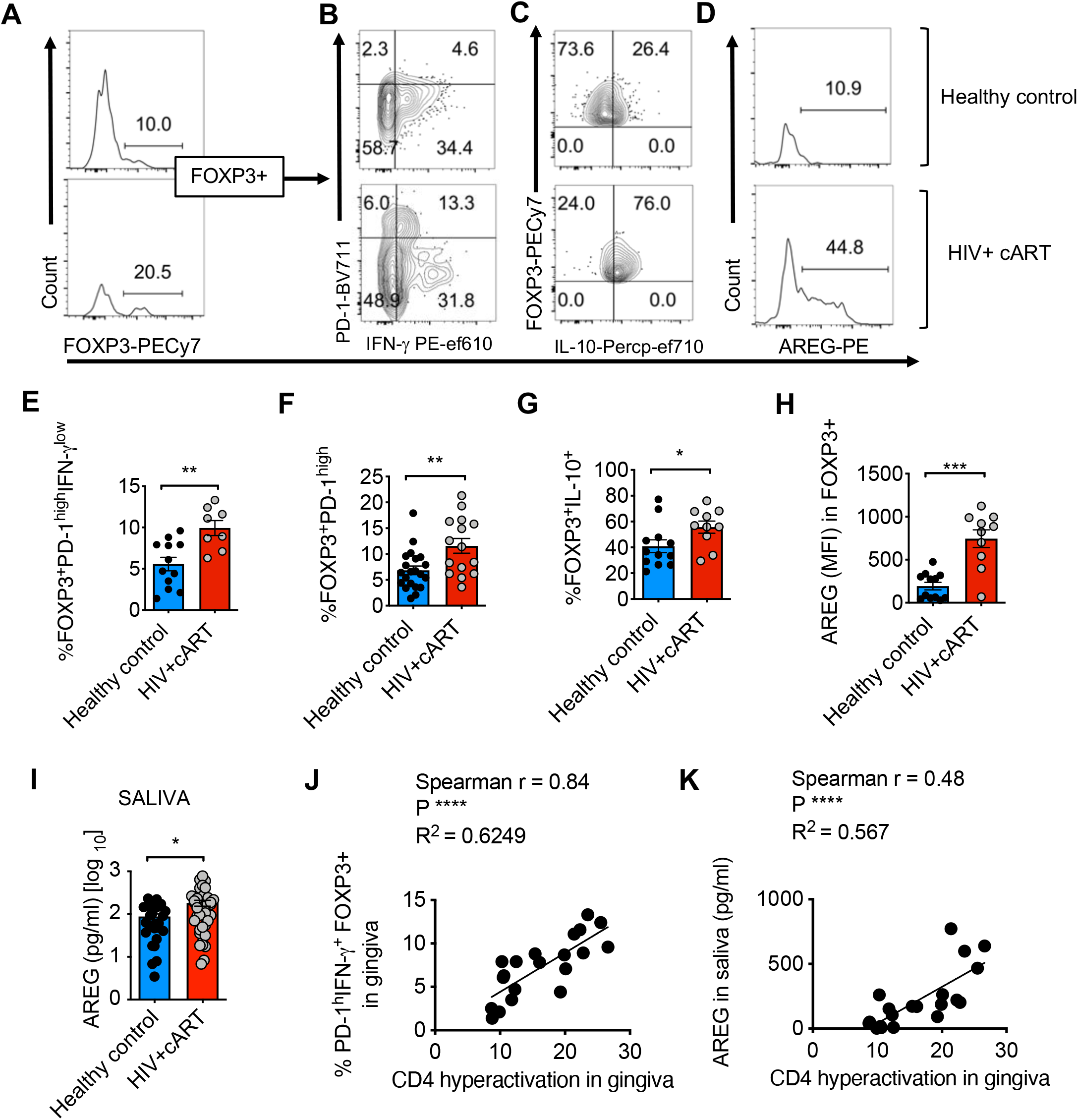
HIV+ patients have an increased abundance of PD-1^hi^CD25^hi^ IFN-γ^+^AREG^hi^ FOXP3^+^ cells correlating with CD4^+^T cell hyperactivation in the oral mucosa. HOILs from gingival mucosa from healthy controls and HIV+ patients on cART were processed for flow cytometry *ex vivo*. (**A**) FOXP3 expression in CD3^+^CD4^+^ gated HOIL cells. PD-1 and IFN-γ (**B**), IL-10(**C**), AREG (**D**), expression in FOXP3^+^ population. Statistical analyses and comparison between the groups for % PD-1^hi^IFN-γ^+^ cells (**E**), % PD-1^hi^ cells (**F**), % IL-10^+^ cells (**G**), and AREG expression (**H**) in FOXP3^+^ population. **I**) ELISA quantification of AREG levels in saliva (* P<0.05; Mann Whitney test). **J, K**) Correlation of % PD-1^hi^CD25^+^ cells in FOXP3^+^ population (**J**) and salivary AREG (**K**), with effector CD4 hyperactivation (% CD38^+^HLADR^+^ in FOXP3^neg^CD4^+^ T cells in gingival mucosa; **Fig.1E**; n = 20).

## Discussion

### A unique population of tissue T_reg_-like FOXP3^+^ cells accumulates in oral MALT and mucosal tissue during HIV infection

Immunological complications in HIV^+^ individuals on treatment appear to result from a self-perpetuating cycle of events involving microbial translocation, excessive release of pro-inflammatory cytokines, and CD4 T cell activation which, in excess, increases the cellular targets for HIV infection and subsequent immune exhaustion^12^. Here we show that HIV and TLR-2 ligands can lead to the accumulation of non-suppressive PD-1^high^ IFN-γ^+^ AREG^+^ FOXP3^+^ cells in an IL-1β dependent manner. These data support the notion that PD-1^+^T_regs_ may intrinsically survive and proliferate better with HIV infection leading to the accumulation of dysfunctional T_regs_. Interestingly a small proportion of PD-1^neg^ T_reg_ or CD45RO^neg^ naive T_reg_ cells can also upregulate PD-1 and IFN-γ in the context of HIV infection (and not TLR-2 stimulation alone; **Fig.5A**, see Uninfected), which also show higher proliferation with TLR-2 and IL-1β stimulation in the context of HIV infection (**Fig. S13, S14**). It is conceivable that in mucosal tissues and tonsils that are known to be enriched with pre-existing PD-1^+^FOXP3^+^ and memory FOXP3^+^ cells, these cells may contribute better to the accrual of dysfunctional T_reg_ cells than naïve T cells. However, the local induction from PD-1^neg^ FOXP3^+^ cells and recently activated FOXP3^+^ cells cannot be ruled out in the process. Therefore we speculate that in HIV^+^ individuals, the effects of HIV, TLR-2 activation, and IL-1β on FOXP3^+^ cell accumulation are driven by complementary and synergistic processes of induction, survival, and proliferation of FOXP3^+^ cells. By demonstrating the mechanistic details by which AKT1 and PD-1 enhance FOXP3 stability and expansion of dysfunctional FOXP3^+^ cells, our study unveils a critical process that may contribute to HIV-mediated CD4^+^ T cell activation that persists in oral mucosa during therapy.

Our study is consistent with previous data showing HIV-1-driven T_reg_ accumulation in lymphoid tissues and the association of TLR-2 ligands and the NLRP3 inflammasome in immune activation and disease progression in HIV/AIDS ^31, 66, 67^. Elevated levels of soluble salivary TLR-2 ligands and CD14 levels in conjunction with higher TLR and inflammasome activation (**Fig.2A, B**) suggest that these features might be strongly linked to the previously established dysbiosis of the oral microbiome in HIV^+^ patients^39^. While the additional role of endocytosed HIV and the resultant TLR-7/8 activation cannot be ruled out ^68^, our data show mechanistic details by which TLR-2 ligands and IL-1β/inflammasome might contribute to proliferation and dysfunction of FOXP3^+^ cells and excessive immune activation in the oral mucosa. The question that remains to be addressed is whether the dysfunction of non-regulatory T cells, and perhaps other cell types, is due to HIV, altered T_reg_ function, or both.

The human tonsil infection model that we employed here has been previously shown to support productive HIV infection, cell death, and release of cytokines such as IL-1β and IL-10, which is similar to an HIV-induced inflammation in humans^34, 50, 69, 70^. Also, being an oral MALT system, it provides a preview into immune cell alterations in oral mucosal tissues. Because the rationale of the study was to determine the underlying oral mucosal dysregulation in HIV+cART patients, we wanted to mimic the effect of HIV in the presence and absence of HIV inhibitor. Considering the different anti-viral regimens that patients take, our *in vitro* experiments may not be exactly physiologically relevant, but may show similarity to HIV+cART patient samples. With this system, we show that FOXP3^+^ cells are highly permissible to HIV infection and undergo cell death during HIV infection (**Fig.3C,4B**). However, a unique population of PD-1^high^IFN-γ^+^AREG^high^FOXP3^+^ cells that expressed anti-apoptotic BCL-2 and are rescued from cell death appeared in cultures, thus unveiling new features of FOXP3^+^ cell dysregulation during HIV infection. These cells expressing CD45RO, IL-1R and ST-2 have an activated/memory T_reg_ like phenotype^63, 71^, and proliferated with TLR-2 agonists and IL-1β even in the presence of HIV reverse transcriptase inhibition (**Fig.5**). These PD-1^hi^ T_regs_ responding to IL-1β and expressing AREG resemble the tissue T_regs_ induced by IL-33, another IL-1 superfamily cytokine. Similar to tissue T_regs_ they may have non-suppressive roles and may function towards mucosal tissue repair during inflammation. They might also differentially govern non-immunological processes in oral mucosa of HIV^+^ patients, compatible with previously described functions of tissue T_reg_ ^57^. PD-1^high^ FOXP3^+^ cells appeared to have low MFI of IFN-γ expression (**Fig.3D, 4C, S16**), consistent with FOXP3 and PD-1 mediated inhibition of IFN-γ^59^. However, these cells from few other experiments showed higher MFI of IFN-γexpression (**Fig.S15**). The reason behind these discordant results is unclear but is likely linked to differences in donors and their underlying tonsil infections. IL-1β enhanced PD-1 upregulation as well as the proliferation of PD-1^high^IFN-γ^+^AREG^high^ FOXP3^+^ cells in a PI-3K/AKT dependent manner(**Fig.5E, S16**). Despite the proliferation and enhanced FOXP3 protein stability conferred by PD-1, these PD-1^high^IFN-γ^+^AREG^high^ FOXP3^+^ cells that are generated during HIV infection lack suppressive ability in blocking CD4 T cell proliferation. This finding concurs with previous data showing cytokines that can activate PI-3K/AKT function or maintain T cell responsiveness to IL-2 in CD4^+^ cells can also abrogate T_reg_ suppression^58–60, 72^. These data are also in accordance with previous studies implicating heightened PD-1 and IFN-γ expression in T_reg_ dysfunction ^73–75^.

BLIMP1 is a transcriptional repressor that is critical for IL-10 expression in TFR cells. TFR cells regulate B cells and TFH cells, thereby controlling the germinal center response, autoantibody production, and autoimmune destruction^53^. Whereas BLIMP1 is expressed in a proportion of lymphoid FOXP3^+^ cells, it is expressed in a majority of FOXP3^+^ cells found in gut mucosa and tissues and is likely crucial for their IL-10 expression in environmental interfaces^76^. Whether the PD-1^high^IFN-γ^+^AREG^high^FOXP3^+^ cells in HIV^+^ oral mucosa that express IL-10 also co-express BLIMP-1 is an important question that remains to be investigated in the future. Although AREG was originally described as an epithelial cell-derived factor and is a member of the epidermal growth factor receptor family, it is now clear that this protein can be expressed by activated immune cells during inflammatory conditions^56^. AREG produced by T cells is critical for type 2 adaptive immune responses and gut epithelial cell proliferation that facilitates helminth parasite clearance. Tissue T_regs_ are shown to express this cytokine and are critical for non-immunological tissue repair functions^57^. Here we show that AREG expression is high in PD-1^high^ FOXP3^+^ cells and is further upregulated by IL-1β in PI-3K/AKT dependent manner. Alarmins such as IL-18 and IL-33 have been previously shown to up-regulate this cytokine in FOXP3^+^ cells^57^. In a tonsillar CD4^+^ T cell environment, although HIV did not induce IL-33 expression (**Fig.S11**), the exogenous addition of both IL-1β and IL-33 upregulated AREG expression in FOXP3^+^cells (**Fig.5D**). However, in the context of HIV infection, only the endogenous IL-1β released due to caspase-1 activity upregulated AREG *in vitro* (**Fig.5B**). PD-1 enhancement and AEP inhibition due to IL-1β upregulates AREG expression in PD-1^high^ FOXP3^+^ cells. The mechanism underlying the inverse relationship between AEP activity and AREG expression remains to be explored in the future.

### *In vivo* evidence for the enrichment of PD-1^high^IFN-γ^+^AREG^high^FOXP3^+^ cells in oral mucosa of HIV+ individuals on therapy

Our results from *in vitro* HIV infection experiments support transcriptomic and flow cytometric profiling results from HIV^+^ patients undergoing suppressive antiviral therapy, whose oral mucosa also revealed TLR signaling pathway upregulation and an inflammasome gene signature that paralleled excessive CD4^+^T cell activation and enrichment of PD-1^high^IFN-γ^+^ AREG^high^FOXP3^+^cells (**Fig.1, 10**). We found distinct populations of CD38^+^HLA-DR^+^ in CD4^+^ T cells only in oral mucosa and not in PBMC (**Fig. 2D**; data not shown). It may be related to the downregulation of HLA-DR (MHC) in CD4^+^ T cells. While HLA-DR downregulation in monocytes is associated with immune-suppression, it remains to be seen if this is the case for CD4^+^ T cells. PD-1^high^IFN-γ^+^ AREG^high^FOXP3^+^cells we describe here resemble the dysfunctional Th1-T_regs_ which also display constitutive activation of PI3K/AKT/Foxo1/3 signaling cascade in multiple sclerosis patients^77^. However, we also show that CD4^+^ T cell hyperactivation and enrichment of PD-1^high^IFN-γ^+^AREG^high^FOXP3^+^cells in oral mucosa of HIV^+^ patients coincide with increased TLR-2 signaling and salivary s-TLR-2 ligands. (**Fig.1B,1D,2B**). We speculate that increased s-TLR-2 ligands we observed in HIV^+^ patients (**Fig.2B**), maybe due to TLR-2 shedding as a consequence of increased pro-inflammatory signaling downstream to the TLR-2 signaling^78^. PD-1 expression on CD4^+^ T cells and T_regs_ is known to be associated with immune activation as well as HIV^+^ reservoirs, and thus this molecule is targeted for therapy in HIV^+^ patients ^74, 79, 80^. Gut mucosa, a tissue enriched with lymphoid structures and bombarded by microbial products as a result of microbial translocation, serves as the largest reservoir^81^. Survival advantage and proliferation of CD4^+^ T cells by homeostatic cytokines and chronic exposure to antigens or other stimulants contribute to the expansion of latently infected cells and consequent viral persistence and establishment of reservoirs ^81^. Therefore, considering that HIV^+^ patients on therapy show oral microbiome dysbiosis^38, 39^, the tissue T_reg_-like PD-1^high^FOXP3^+^ cells appear to fit these criteria and might provide a supportive environment for the maintenance of HIV reservoirs in the oral mucosa. This tenet is consistent with previous reports showing that FOXP3^+^ cells are highly permissible and contribute to latent reservoir compartments^82, 83^. Future studies are required to conclusively verify this possibility in the oral mucosal environment. PD-1^high^IFN-γ^+^ AREG^high^FOXP3^+^cells could not suppress CD4^+^ T cells *in vitro*. These T_reg_ cells with high expression of BCL-2 and KI-67, and resistance to apoptosis (Fig.3E, **4D**), suggest that they may be long-lived and may undergo continuous cycling. This is consistent to previous reports showing T_regs_’ resistance to apoptosis^84, 85^. Here we also show that these FOXP3^+^ cells are dysfunctional and have a unique phenotype. Moreover, increases in PD-1^high^IFN-γ^+^AREG^high^FOXP3^+^cells correlating with CD4 hyperactivation in the oral mucosa of HIV+cART patients (**Fig.8**), would imply that these may be long-lived and dysfunctional in these patients. However, we cannot rule out that these cells co-expressing IL-10 may still inhibit myeloid populations, neutrophils, and resident macrophages, providing an immune-suppressive environment. Taken together, these results show that persistent microbial stimulants and excessive IL-1β signaling perturb FOXP3^+^ T cell homeostasis and function, and underlie the processes of residual oral mucosal immune dysfunction in HIV^+^ patients on therapy.

## Author contributions

PP designed the study, performed experiments, analyzed data, supervised the project, and wrote the manuscript. FF and AP provided gingival biopsies from human participants, and RA referred the patients to the study. NB and ES performed the experiments, analyzed ELISA data, and contributed to discussions. ES obtained consents from patients, collected the saliva and blood, and performed ELISA. AT provided statistical analysis consultation for bioinformatics data. ADL, NG, JK, and ML read the manuscript and contributed to discussions.

## Acknowledgments

PP was supported by CWRU Center for AIDS Research (CFAR) Catalytic award, and RO1DE026923 NIH/NIDCR funding. We thank Jennifer Bongorno-Hurt for preparing HIV-viral stocks and Sangeetha Jayaraman for technical assistance with patient recruitment paperwork and assessing data in a masked fashion. We acknowledge Rafick P Sekaly for critically reading the manuscript and valuable suggestions. We thank Ms. Patricia Mehosky for proof-reading the manuscript.

## Methods

### Human PBMC, gingival biopsies, and tonsils

Human blood, gingival biopsies, and saliva were obtained with informed consents from healthy individuals and Cleveland HIV^+^ cohort under a protocol approved by the University Hospitals the Cleveland Medical Center Institutional Review Board. Healthy control subjects were at least 18 years of age and in good general health (**Table.1**). Exclusion criteria were oral inflammatory lesions (including gingivitis and periodontitis), oral cancer diagnosis, soft tissue lesions, and the use of tobacco in the past month. HIV^+^ participants were 18 years or older, and were HIV positive with cART treatment for at least 1 year. > 75 % of HIV+ patients reported prior and current soft tissue lesions, gingivitis, and periodontitis. Exclusion criteria were oral cancer diagnosis and the use of tobacco in the past month. The absence of tobacco use was confirmed by Cotinine ELISA in saliva. CD4^+^ counts were at least 350 - 700/μl for the control and HIV^+^ patients. Palatine tonsils were obtained as discards from tonsillectomy surgeries performed at University Hospitals Cleveland Medical Center through the Histology Tissue Procurement Facility following an IRB-approved protocol. PBMC were collected from blood using Ficoll-Paque PLUS centrifugation and subsequent washing with PBS. A single-cell suspension of gingival tissues and tonsils were prepared by Collagenase 1A digestion and processed for flow cytometry or cell culture.

### HIV infection *in vitro*

HIV infections in tonsil cultures were performed using X4-tropic NL43-GFP-IRES-Nef or HIV-NLGNef, a recombinant virus with NL4-3 backbone expressing Green Fluorescent Protein (GFP) and Nef on a bicistronic transcript^47, 86^. Viral constructs were obtained through NIH AIDS Reagent Program and the viruses were generated by transfecting 293 T cells with proviral DNA. The R5-tropic virus was created replacing the Env in NL43-GFP-IRES-Nef with the EcoR1-Bam fragment from NLAD8, an NL43 construct containing CCR5-tropic HIV-1 ADA envelope ^47, 48^. Concentrated virus stock titers were determined by p24 enzyme-linked immunosorbent assay (ELISA). For infections, tonsil tissues were digested using collagenase, and a single-cell suspension of the human tonsillar culture (HTC) (1 million cells/ well) was plated with α-CD3 (1μg/ml) and α-CD28 (1 μg/ml) TCR activating antibodies in U-bottom 96 well plates at least in triplicate wells. After 48 hours, the bulk HTC were spinoculated with replication-competent HIV-1 NLAD8-GFP virus stock (70 ng of p24/10^6^ cells). Cells were rested for 48 hours in medium without TCR activation in select experiments. As indicated in some experiments, purified CD4^+^ T cells and T_regs_ were used instead of whole HTC. 24-36 hours post-infection, 50% of the cells and media were removed and replaced with media containing fresh media, Efavirenz, and indicated cytokines or reagents. This allowed an initial round of infection and cell death to occur before the addition of the indicated reagents. When indicated, TGF-β1 (10 ng/ml) and IL-2 (100 U/ml) were also added during this time to induce and maintain FOXP3^+^ cells. Confirmatory experiments were performed using both X4- and R5-tropic viruses^86^. Cells were cultured in complete RPMI-1640 (Hyclone) supplemented with 10% human serum, 100 U/ml penicillin, 100 µg/ml streptomycin, 2 mM glutamine, 10 mM HEPES and 1 mM sodium pyruvate. To determine productive HIV infection (GFP) and regulation of protein expression, cells were analyzed by flow cytometry on day 2 – day 8 post-infection. Flow cytometry analyses and ELISAs were performed in triplicates using tonsils from at least three independent donors.

### Antibodies and reagents

Unconjugated or fluorochrome-conjugated antibodies for human CD28(CD28.2), CD25 (M-A251), CD4 (OKT4), CD45(HI30), CD8 (RPA-T8), HLA-DR(LN3), IFN-γ (4S.B3), IL-17A(eBio64DEC17), FOXP3(236A/E7), Phospho-AKT 1 (Ser473)(SDRNR), BCL-6 (BCL-UP), CXCR5(MU5UBEE), Ki-67 (SolA15), IL-10 (JES3-9D7), AREG (AREG559), IL-6(MQ2-13A5), ST2 (goat polyclonal), phospho-caspase 1 (polyclonal), LY294002, and Cell-trace violet were all purchased from Thermofisher Scientific. CD279 (PD-1)(EH12.1), CXCR4(12g5), CCR5 (2D7/CCR5), BCL-2(Bcl-2/100), CD19 (SJ25C1), CD38 (HIT2), CD3 (HIT3a), and IL-1R1(hIL1R-M1) were from BD Biosciences. Phospho-AEP(SER 226) antibody, Efavirenz (SML1284-1ML), and AEP inhibitor were from Millipore Sigma. Biotinylated antibody for AEP, BLIMP1 antibody(646702), Human TGF-β1, and the chimeric PDL-1-Fc were purchased from R&D systems. Appropriate secondary antibodies such as secondary donkey anti-mouse IgG-BV421 (for IL1-RI staining), anti-goat IgG (H+L) superclonal™-Alexa Fluor 647 (for ST2 staining) streptavidin-APC, and anti-rabbit PE or APC antibodies were purchased from Jackson Immunoresearch. Anti-Biotin multi-sort and human CD4+CD45RO+ isolation kits were purchased from Miltenyi Biotec (Auburn, CA). PE+ cell, CD4+ T cell, and T_reg_ isolation kits were purchased from Stem Cell Technologies (Vancouver, Canada). Recombinant IL-2, IL-1β, and IL-33 cytokines were purchased from BioBasic Inc. (Amherst, NY). s-CD14, s-TLR-2, Cotinine, AREG, IL-1β, and IL-6 ELISA kits were from Boster Bio (Pleasanton, CA). IL-1 receptor antagonist Anakinra was a kind gift from Dr. Su at NIAID, NIH. Nigericin and PG-LPS were purchased from Invivogen. Heat killed *Candida albicans* germ tubes (HKGT) were prepared in the laboratory by growing the blastospores (10*9 / ml) into germ-tubes in complete RPMI at 37°C for 4-6 hours, and heat killing the germ tubes at 75°C for 60 minutes.

### Fluorochrome antibody staining and flow cytometry

For single-cell flow cytometry analyses, surface receptors were first stained using the antibodies in PBS/BSA. Live-Dead viability staining was performed to detect and remove dead cells in the analyses. For FOXP3 and other intracellular protein stainings, the cells were fixed with a FOXP3 fixation-permeabilization set (Thermofisher Scientific) after the surface staining. Unstimulated-, un-stain-, isotype-, secondary antibody alone-, single stain-, and FMO-controls were included in all the preliminary and confirmatory experiments, and appropriate controls were chosen. Before intracellular cytokine staining, cultures were re-stimulated with PMA (50 ng/ml) and Ionomycin (500 ng/ml) for 4 hours, with brefeldin-A (10 µg/ml) added in the last 2 hours. For p-AKT1 staining, the cells were washed, fixed, and stained with a Phosflow staining kit (BD Biosciences) using the manufacturer’s protocol. Data were acquired using BD Fortessa cytometers and analyzed using FlowJo 9.8 or 10.5.3 software. Populations were pre-gated for lymphocyte, singlet, viable, CD3^+^, and CD8^-^ or CD4^+^ cells during flow cytometry analyses, unless otherwise specified.

### PD-1 engagement and T_reg_ suppression assay *in vitro*

Tonsil cells were stimulated and infected in U-bottom 96 well plates as above. 36 hours after infection, the cells were moved to the plate coated with PDL-1-Fc for PD-1 engagement. The plates were previously coated with PDL-1-Fc for 12-16 hours. Appropriate isotype control (IgG2a) was used in control wells. Flow cytometry was performed on day 4 or 5 after PD-1 engagement. For the suppression assay, three groups of magnetic sorted cells purified *ex vivo* from tonsils were activated with CD3/CD28 antibodies with added TGF-β1 and IL-2 for 96 hours; I) CD45RO^neg^ naïve CD4^+^CD25^+^CD127^low^T_regs_ (>90% FOXP3^+^), II) Purified CD4^+^CD25^-^T cells that were subsequently used as responder T cells (T_resp_) in the co-culture assay and III) Purified CD4^+^ T cells infected with HIV. PD-1^high^CD25^+^ cells purified from these cultures were 80-88% FOXP3^+^, IFN-γ^+^(52%) (**Fig.S18**), AREG^high^, and were used as PD-1^high^ CD25^+^FOXP3^+^ cells. For co-culture T_reg_ suppression assay, 3 x 10^4^ T_resp_ cells were labeled with cell-trace violet, and co-cultured with 3 x 10^4^ CD4^+^CD25^+^CD127^low^T_regs_, or 3 x 10^4^ PD-1^high^ CD25^+^ cells in triplicate wells of U bottomed 96-well plates in the presence of soluble 1 μg/mlα-CD3 and 1 μg/ml α-CD28 antibodies for the indicated duration^58^.

### RNA sequencing

Sample preparation, sequencing, alignment, and data analyses were performed by Novogene genomic services. Strand-specific whole transcriptome sequencing libraries were prepared using NEB Next® Ultra™ RNA Library Prep Kit. The sequencing used a paired-end protocol (PE150). Indexed RNA-seq libraries were sequenced on a HiSeq2500 with Illumina TruSeq V4 chemistry (Illumina, San Diego, CA, USA). The FASTQ files with 125bp paired-end reads were processed using Trimmomatic (version 0.30) to remove adaptor sequences. The trimmed FASTQ data were aligned to the human genome with STAR (version 2.4.2a), which used GENCODE gtf file version 4 (Ensembl 78). Differential expression analysis: The gene reads count data from HOIL and PBMC samples, each derived from three independent human individuals were normalized with R Package limma (version 3.26.8) and analyzed with an unpaired t-test. HOIL samples from three control individuals were pooled and compared with three independent HIV^+^ individuals. The normalized reads count data were used to generate RPKM values for the heatmap display. Pathway analysis and heat maps: The Differential expressed gene list (DEG) was generated using unbiased molecular and cellular functional analyses. Heatmaps for different cytokine signatures were created in R using the heatmap.2 function in g plots (version 2.17.0). Gene set enrich analysis (GSEA) was performed using the GSEA software obtained from the Broad Institute (http://www.broad.mit.edu/GSEA). REACTOME, GO and MSigDB gene sets and reference pathways were employed when relevant. The whole gene list was ranked before uploading to the GSEA software for pathway analysis. Normalized maximum deviation from zero was recorded as the enrichment score and normalized for obtaining normalized enrichment score (NES).

### Statistical analyses

P values were calculated by Mann-Whitney test in Prism 8 (GraphPad Software, Inc.) assuming random distribution unless otherwise noted. For some multiple comparisons within *in vitro* culture groups, one-way ANOVA was used. Unpaired t-test and two-way ANOVA were used for multiple comparisons between two or more groups. Bonferroni t-test was the post hoc test used for multiple comparisons. *P < 0.05 were considered significant. To measure the strength of the association, correlation plots, spearman (r), simple linear regression analyses (R^2^) were used, and an alpha value of *<0.05 was considered significant.

## Supplementary materials

Table and supplementary figures are provided as supplementary material.

## References

1. Ferreira, S. et al. Prevalence of oral manifestations of HIV infection in Rio De Janeiro, Brazil from 1988 to 2004. AIDS patient care and STDs 21, 724–731 (2007).

2. Patton, L.L. Oral lesions associated with human immunodeficiency virus disease. Dental clinics of North America 57, 673–698 (2013).

3. Shiboski, C.H. et al. Overview of the oral HIV/AIDS Research Alliance Program. Advances in dental research 23, 28–33 (2011).

4. Webster-Cyriaque, J., Duus, K., Cooper, C. & Duncan, M. Oral EBV and KSHV infection in HIV. Advances in dental research 19, 91–95 (2006).

5. Vernon, L.T. et al. Effect of Nadir CD4+ T cell count on clinical measures of periodontal disease in HIV+ adults before and during immune reconstitution on HAART. PloS one 8, e76986 (2013).

6. Vernon, L.T. et al. Comorbidities associated with HIV and antiretroviral therapy (clinical sciences): a workshop report. Oral diseases 22 Suppl 1, 135–148 (2016).

7. Mendez-Lagares, G., Leal, M., del Pozo-Balado, M.M., Leon, J.A. & Pacheco, Y.M. Is it age or HIV that drives the regulatory T-cells expansion that occurs in older HIV-infected persons? Clinical immunology 136, 157–159; author reply 160 (2010).

8. Tappuni, A.R. Immune reconstitution inflammatory syndrome. Advances in dental research 23, 90–96 (2011).

9. Tenorio, A.R., Martinson, J., Pollard, D., Baum, L. & Landay, A. The relationship of T-regulatory cell subsets to disease stage, immune activation, and pathogen-specific immunity in HIV infection. Journal of acquired immune deficiency syndromes 48, 577–580 (2008).

10. Caruso, M.P. et al. Impact of HIV-ART on the restoration of Th17 and Treg cells in blood and female genital mucosa. Scientific reports 9, 1978 (2019).

11. Rueda, C.M., Velilla, P.A., Chougnet, C.A., Montoya, C.J. & Rugeles, M.T. HIV-induced T-cell activation/exhaustion in rectal mucosa is controlled only partially by antiretroviral treatment. PloS one 7, e30307 (2012).

12. Gallo, R.C. HIV/AIDS Research for the Future. Cell host & microbe 27, 499–501 (2020).

13. Catalfamo, M., Le Saout, C. & Lane, H.C. The role of cytokines in the pathogenesis and treatment of HIV infection. Cytokine & growth factor reviews 23, 207–214 (2012).

14. Younes, S.A. et al. Cycling CD4+ T cells in HIV-infected immune nonresponders have mitochondrial dysfunction. The Journal of clinical investigation 128, 5083–5094 (2018).

15. El Howati, A. & Tappuni, A. Systematic review of the changing pattern of the oral manifestations of HIV. J Investig Clin Dent 9, e12351 (2018).

16. Sokoya, T., Steel, H.C., Nieuwoudt, M. & Rossouw, T.M. HIV as a Cause of Immune Activation and Immunosenescence. Mediators Inflamm 2017, 6825493 (2017).

17. Heron, S.E. & Elahi, S. HIV Infection and Compromised Mucosal Immunity: Oral Manifestations and Systemic Inflammation. Frontiers in immunology 8, 241 (2017).

18. Weinberg, A. et al. Innate immune mechanisms to oral pathogens in oral mucosa of HIV-infected individuals. Oral diseases 26 Suppl 1, 69–79 (2020).

19. Gaffen, S.L. & Moutsopoulos, N.M. Regulation of host-microbe interactions at oral mucosal barriers by type 17 immunity. Sci Immunol 5 (2020).

20. Dutzan, N. et al. A dysbiotic microbiome triggers TH17 cells to mediate oral mucosal immunopathology in mice and humans. Sci Transl Med 10 (2018).

21. Sultan, A.S., Kong, E.F., Rizk, A.M. & Jabra-Rizk, M.A. The oral microbiome: A Lesson in coexistence. PLoS Pathog 14, e1006719 (2018).

22. George, J., Wagner, W., Lewis, M.G. & Mattapallil, J.J. Significant Depletion of CD4(+) T Cells Occurs in the Oral Mucosa during Simian Immunodeficiency Virus Infection with the Infected CD4(+) T Cell Reservoir Continuing to Persist in the Oral Mucosa during Antiretroviral Therapy. Journal of immunology research 2015, 673815 (2015).

23. Oswald-Richter, K. et al. HIV infection of naturally occurring and genetically reprogrammed human regulatory T-cells. PLoS Biol 2, E198 (2004).

24. Chevalier, M.F. et al. The Th17/Treg ratio, IL-1RA and sCD14 levels in primary HIV infection predict the T-cell activation set point in the absence of systemic microbial translocation. PLoS pathogens 9, e1003453 (2013).

25. Raynor, J. et al. IL-6 and ICOS Antagonize Bim and Promote Regulatory T Cell Accrual with Age. J Immunol 195, 944–952 (2015).

26. Kanwar, B., Favre, D. & McCune, J.M. Th17 and regulatory T cells: implications for AIDS pathogenesis. Current opinion in HIV and AIDS 5, 151–157 (2010).

27. Holmes, D., Jiang, Q., Zhang, L. & Su, L. Foxp3 and Treg cells in HIV-1 infection and immuno-pathogenesis. Immunologic research 41, 248–266 (2008).

28. Moreno-Fernandez, M.E., Presicce, P. & Chougnet, C.A. Homeostasis and function of regulatory T cells in HIV/SIV infection. Journal of virology 86, 10262–10269 (2012).

29. Epple, H.J. et al. Mucosal but not peripheral FOXP3+ regulatory T cells are highly increased in untreated HIV infection and normalize after suppressive HAART. Blood 108, 3072–3078 (2006).

30. Eggena, M.P. et al. Depletion of regulatory T cells in HIV infection is associated with immune activation. Journal of immunology 174, 4407–4414 (2005).

31. Nilsson, J. et al. HIV-1-driven regulatory T-cell accumulation in lymphoid tissues is associated with disease progression in HIV/AIDS. Blood 108, 3808–3817 (2006).

32. Estes, J.D. et al. Simian immunodeficiency virus-induced lymphatic tissue fibrosis is mediated by transforming growth factor beta 1-positive regulatory T cells and begins in early infection. The Journal of infectious diseases 195, 551–561 (2007).

33. Kinter, A.L. et al. CD25(+)CD4(+) regulatory T cells from the peripheral blood of asymptomatic HIV-infected individuals regulate CD4(+) and CD8(+) HIV-specific T cell immune responses in vitro and are associated with favorable clinical markers of disease status. The Journal of experimental medicine 200, 331–343 (2004).

34. Chevalier, M.F. et al. Phenotype alterations in regulatory T-cell subsets in primary HIV infection and identification of Tr1-like cells as the main interleukin 10-producing CD4+ T cells. The Journal of infectious diseases 211, 769–779 (2015).

35. Subramanian, A. et al. Gene set enrichment analysis: a knowledge-based approach for interpreting genome-wide expression profiles. Proceedings of the National Academy of Sciences of the United States of America 102, 15545–15550 (2005).

36. Lederman, M.M. et al. Immunologic failure despite suppressive antiretroviral therapy is related to activation and turnover of memory CD4 cells. The Journal of infectious diseases 204, 1217–1226 (2011).

37. Shive, C.L. et al. Inflammatory cytokines drive CD4+ T-cell cycling and impaired responsiveness to interleukin 7: implications for immune failure in HIV disease. The Journal of infectious diseases 210, 619–629 (2014).

38. Mukherjee, P.K. et al. Dysbiosis in the oral bacterial and fungal microbiome of HIV-infected subjects is associated with clinical and immunologic variables of HIV infection. PloS one 13, e0200285 (2018).

39. Presti, R.M. et al. Alterations in the oral microbiome in HIV-infected participants after antiretroviral therapy administration are influenced by immune status. Aids 32, 1279–1287 (2018).

40. Pandiyan, P. et al. Microbiome dependent regulation of Tregs and Th17 cells in mucosa. Front Immunol 3 (2019).

41. Bhaskaran, N. et al. Role of Short Chain Fatty Acids in Controlling Tregs and Immunopathology During Mucosal Infection. Front Microbiol 9, 1995 (2018).

42. Bhaskaran, N., Cohen, S., Zhang, Y., Weinberg, A. & Pandiyan, P. TLR-2 Signaling Promotes IL-17A Production in CD4+CD25+Foxp3+ Regulatory Cells during Oropharyngeal Candidiasis. Pathogens 4, 90–110 (2015).

43. Round, J.L. & Mazmanian, S.K. Inducible Foxp3+ regulatory T-cell development by a commensal bacterium of the intestinal microbiota. Proc Natl Acad Sci U S A 107, 12204–12209 (2010).

44. Dutzan, N., Konkel, J.E., Greenwell-Wild, T. & Moutsopoulos, N.M. Characterization of the human immune cell network at the gingival barrier. Mucosal Immunol 9, 1163–1172 (2016).

45. Miller, S.M. et al. Follicular Regulatory T Cells Are Highly Permissive to R5-Tropic HIV-1. Journal of virology 91 (2017).

46. Banga, R. et al. PD-1(+) and follicular helper T cells are responsible for persistent HIV-1 transcription in treated aviremic individuals. Nature medicine 22, 754–761 (2016).

47. Levy, D.N., Aldrovandi, G.M., Kutsch, O. & Shaw, G.M. Dynamics of HIV-1 recombination in its natural target cells. Proceedings of the National Academy of Sciences of the United States of America 101, 4204–4209 (2004).

48. Cenker, J.J., Stultz, R.D. & McDonald, D. Brain Microglial Cells Are Highly Susceptible to HIV-1 Infection and Spread. AIDS research and human retroviruses 33, 1155–1165 (2017).

49. Jiang, Q. et al. FoxP3+CD4+ regulatory T cells play an important role in acute HIV-1 infection in humanized Rag2-/-gammaC-/- mice in vivo. Blood 112, 2858–2868 (2008).

50. Doitsh, G. et al. Cell death by pyroptosis drives CD4 T-cell depletion in HIV-1 infection. Nature 505, 509–514 (2014).

51. Pan, D. et al. Lack of interleukin-10-mediated anti-inflammatory signals and upregulated interferon gamma production are linked to increased intestinal epithelial cell apoptosis in pathogenic simian immunodeficiency virus infection. Journal of virology 88, 13015–13028 (2014).

52. McCune, J.M. The dynamics of CD4+ T-cell depletion in HIV disease. Nature 410, 974–979 (2001).

53. Cretney, E., Kallies, A. & Nutt, S.L. Differentiation and function of Foxp3(+) effector regulatory T cells. Trends in immunology 34, 74–80 (2013).

54. Gonzalez, O.A., Li, M., Ebersole, J.L. & Huang, C.B. HIV-1 reactivation induced by the periodontal pathogens Fusobacterium nucleatum and Porphyromonas gingivalis involves Toll-like receptor 2 [corrected] and 9 activation in monocytes/macrophages. Clinical and vaccine immunology : CVI 17, 1417–1427 (2010).

55. Mataftsi, M., Skoura, L. & Sakellari, D. HIV infection and periodontal diseases: an overview of the post-HAART era. Oral diseases 17, 13–25 (2011).

56. Zaiss, D.M.W., Gause, W.C., Osborne, L.C. & Artis, D. Emerging functions of amphiregulin in orchestrating immunity, inflammation, and tissue repair. Immunity 42, 216–226 (2015).

57. Arpaia, N. et al. A Distinct Function of Regulatory T Cells in Tissue Protection. Cell 162, 1078–1089 (2015).

58. Pandiyan, P., Zheng, L., Ishihara, S., Reed, J. & Lenardo, M.J. CD4(+)CD25(+)Foxp3(+) regulatory T cells induce cytokine deprivation-mediated apoptosis of effector CD4(+) T cells. Nat Immunol 8, 1353–1362 (2007).

59. Pandiyan, P. & Zhu, J. Origin and functions of pro-inflammatory cytokine producing Foxp3(+) regulatory T cells. Cytokine 76, 13–24 (2015).

60. Ben-Sasson, S.Z., Wang, K., Cohen, J. & Paul, W.E. IL-1beta strikingly enhances antigen-driven CD4 and CD8 T-cell responses. Cold Spring Harb Symp Quant Biol 78, 117–124 (2013).

61. Jain, A., Song, R., Wakeland, E.K. & Pasare, C. T cell-intrinsic IL-1R signaling licenses effector cytokine production by memory CD4 T cells. Nat Commun 9, 3185 (2018).

62. Luo, C.T., Liao, W., Dadi, S., Toure, A. & Li, M.O. Graded Foxo1 activity in Treg cells differentiates tumour immunity from spontaneous autoimmunity. Nature 529, 532–536 (2016).

63. DiSpirito, J.R. et al. Molecular diversification of regulatory T cells in nonlymphoid tissues. Sci Immunol 3 (2018).

64. Manoury, B. et al. An asparaginyl endopeptidase processes a microbial antigen for class II MHC presentation. Nature 396, 695–699 (1998).

65. Stathopoulou, C. et al. PD-1 Inhibitory Receptor Downregulates Asparaginyl Endopeptidase and Maintains Foxp3 Transcription Factor Stability in Induced Regulatory T Cells. Immunity 49, 247–263 e247 (2018).

66. Bandera, A. et al. The NLRP3 Inflammasome Is Upregulated in HIV-Infected Antiretroviral Therapy-Treated Individuals with Defective Immune Recovery. Frontiers in immunology 9, 214 (2018).

67. Tan, D.B. et al. TLR2-induced cytokine responses may characterize HIV-infected patients experiencing mycobacterial immune restoration disease. Aids 25, 1455–1460 (2011).

68. Meas, H.Z. et al. Sensing of HIV-1 by TLR8 activates human T cells and reverses latency. Nat Commun 11, 147 (2020).

69. Andersson, J. et al. The prevalence of regulatory T cells in lymphoid tissue is correlated with viral load in HIV-infected patients. Journal of immunology 174, 3143–3147 (2005).

70. Soare, A.Y. et al. P2X Antagonists Inhibit HIV-1 Productive Infection and Inflammatory Cytokines Interleukin-10 (IL-10) and IL-1beta in a Human Tonsil Explant Model. Journal of virology 93 (2019).

71. Raffin, C., Raimbaud, I., Valmori, D. & Ayyoub, M. Ex vivo IL-1 receptor type I expression in human CD4+ T cells identifies an early intermediate in the differentiation of Th17 from FOXP3+ naive regulatory T cells. J Immunol 187, 5196–5202 (2011).

72. Pandiyan, P., Zheng, L. & Lenardo, M.J. The molecular mechanisms of regulatory T cell immunosuppression. Frontiers in immunology 2, 60 (2011).

73. Lowther, D.E. et al. PD-1 marks dysfunctional regulatory T cells in malignant gliomas. JCI Insight 1 (2016).

74. Xiao, J. et al. PD-1 Upregulation Is Associated with Exhaustion of Regulatory T Cells and Reflects Immune Activation in HIV-1-Infected Individuals. AIDS research and human retroviruses 35, 444–452 (2019).

75. Overacre-Delgoffe, A.E. et al. Interferon-gamma Drives Treg Fragility to Promote Anti-tumor Immunity. Cell 169, 1130–1141 e1111 (2017).

76. Rubtsov, Y.P. et al. Regulatory T cell-derived interleukin-10 limits inflammation at environmental interfaces. Immunity 28, 546–558 (2008).

77. Kitz, A. et al. AKT isoforms modulate Th1-like Treg generation and function in human autoimmune disease. EMBO reports 17, 1169–1183 (2016).

78. Langjahr, P. et al. Metalloproteinase-dependent TLR2 ectodomain shedding is involved in soluble toll-like receptor 2 (sTLR2) production. PloS one 9, e104624 (2014).

79. Fromentin, R. et al. PD-1 blockade potentiates HIV latency reversal ex vivo in CD4(+) T cells from ART-suppressed individuals. Nat Commun 10, 814 (2019).

80. Swindells, S. & Landay, A.L. Time to Study Immune Checkpoint Inhibitors in Patients With HIV Infection and Cancer. JCO Oncol Pract 16, 327–328 (2020).

81. Cohn, L.B., Chomont, N. & Deeks, S.G. The Biology of the HIV-1 Latent Reservoir and Implications for Cure Strategies. Cell host & microbe 27, 519–530 (2020).

82. Tran, T.A. et al. Resting regulatory CD4 T cells: a site of HIV persistence in patients on long-term effective antiretroviral therapy. PloS one 3, e3305 (2008).

83. Li, G. et al. Regulatory T Cells Contribute to HIV-1 Reservoir Persistence in CD4+ T Cells Through Cyclic Adenosine Monophosphate-Dependent Mechanisms in Humanized Mice In Vivo. The Journal of infectious diseases 216, 1579–1591 (2017).

84. Bhaskaran, N. et al. Transforming growth factor-beta1 sustains the survival of Foxp3(+) regulatory cells during late phase of oropharyngeal candidiasis infection. Mucosal immunology 9, 1015–1026 (2016).

85. Chevalier, M.F. & Weiss, L. The split personality of regulatory T cells in HIV infection. Blood 121, 29–37 (2013).

86. Reyes-Rodriguez, A.L., Reuter, M.A. & McDonald, D. Dendritic Cells Enhance HIV Infection of Memory CD4(+) T Cells in Human Lymphoid Tissues. AIDS research and human retroviruses 32, 203–210 (2016).

