## Supplementary figures for "PD-1 dependent expansion of Amphregulin^+^FOXP3^+^ cells is associated with oral immune dysfunction in HIV patients on therapy"

### Supplementary materials

**Table: 1. Human participants enrolled in the study**

| Group | HIV− (n = 32) | HIV+ ART+ (n=46) |
|---|---|---|
| Age (years) median | 49 +/- 9.9 | 54 +/- 18.5 |
| Aged 60 and above | 21.8% | 22.2% |
| Time under cART median | 0 | 15 +/- 7.6 yrs |
| Viral load median | 0 | 20 (range 0.8 – 272) |
| % prior Candidiasis positive | 0 | 34.9% |
| % periodontitis + | 0 | 4.7% |

**Table 1**

Fig. S1

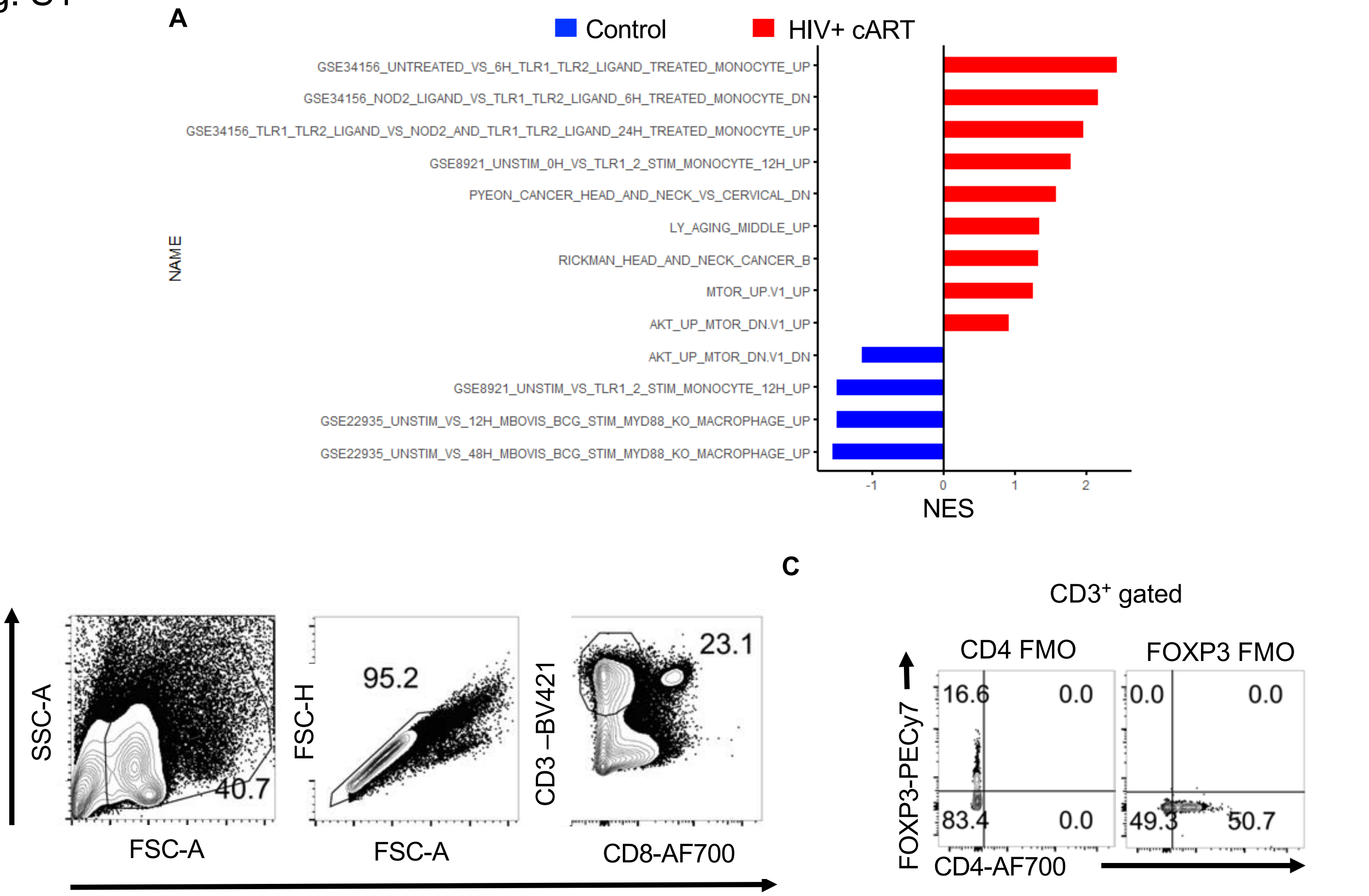


**Fig.S1. Oral Mucosa characterization.** A) Gingival cells were enriched for immune cells by reducing the epithelial cells through gradient centrifugation before transcriptome analyses (n=3 HIV+cART; n=3 uninfected healthy controls). Normalized Enrichment Score (NES) showing the enrichment of pathways of aging, head and cancer and Akt signaling in oral mucosa of HIV$^+$ patients based on gene sets in GO pathways and MSigDB. HOIL were processed *ex vivo* for flow cytometry. CD4$^+$ T cells were gated by either gating on CD8 negative CD3$^+$ cells, or CD4$^+$CD3$^+$ cells. CD8-negative gating approach (B), and FMO controls for CD4 and FOXP3 staining (C) are shown.

Fig.S2

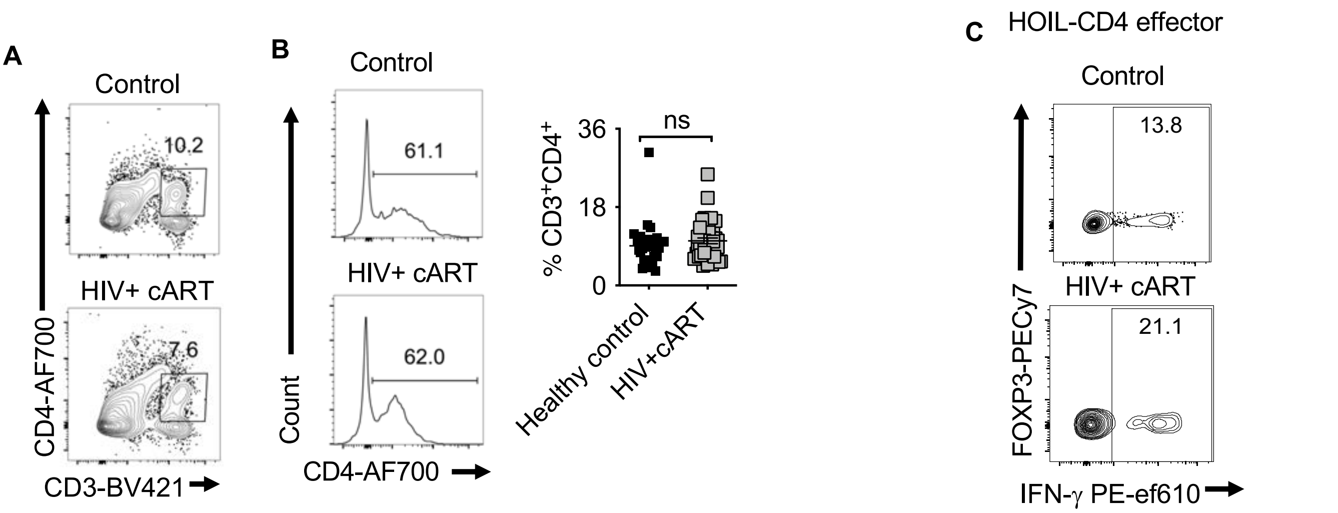


**Fig.S2. Oral Mucosa characterization.** HOIL from study participants (n=46 HIV+ cART; n=32 uninfected healthy controls) were processed *ex vivo*. CD4 cell gating (A), Histogram plots and statistical analyses for CD4 expression in CD3 population (B) and IFN-γ expression in CD4$^+$ FOXP3 negative T cells (C).

Fig.S3

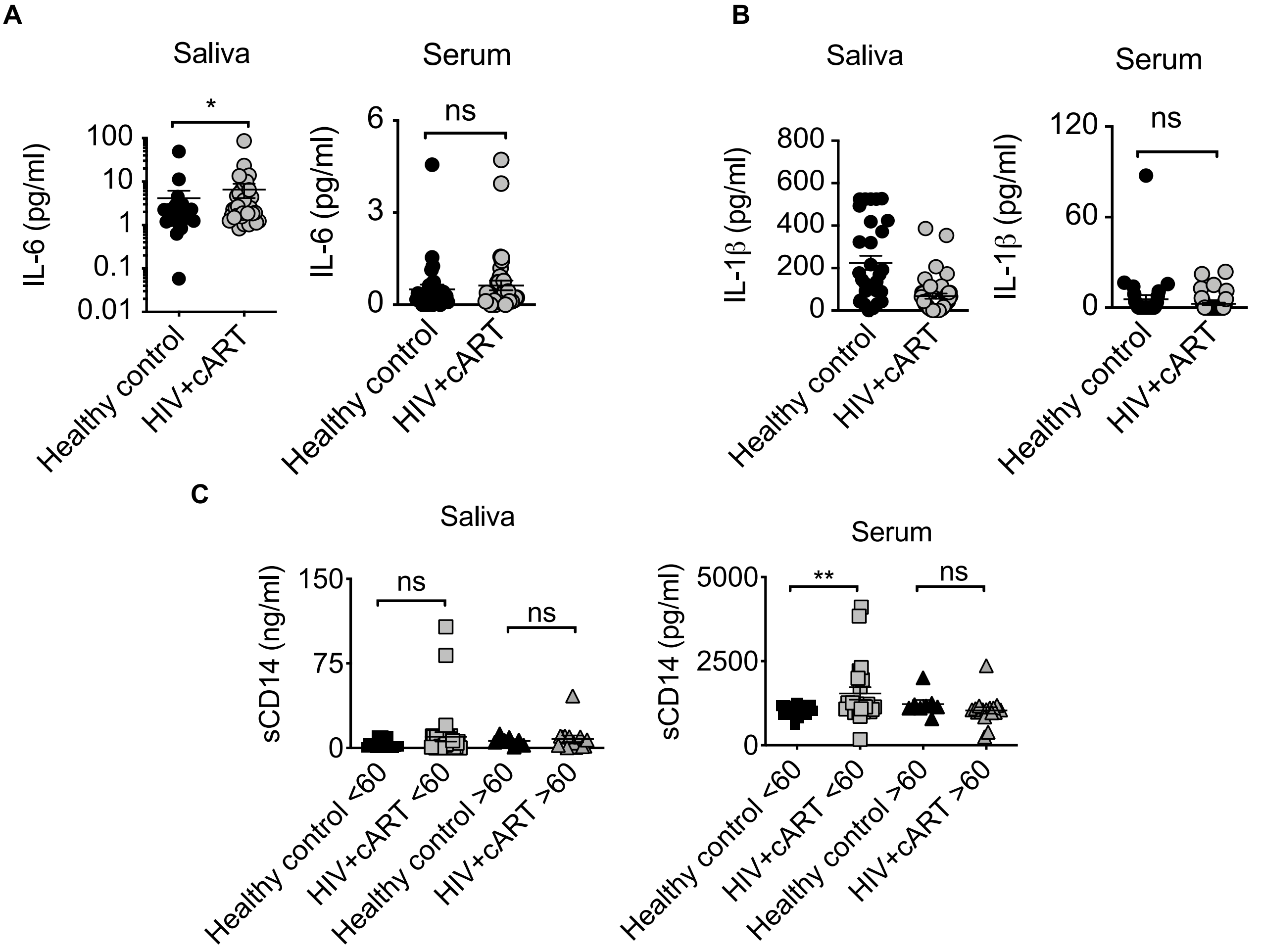


**Fig.S3. Comparison of saliva and serum of HIV+ cART patients.** Saliva and serum from study participants (n=46 HIV+ cART; n=32 uninfected healthy controls) were processed for ELISA. ELISA quantification of IL-6 (A) and IL-1β (B) and soluble CD14 (C) in saliva and serum.

Fig.S4

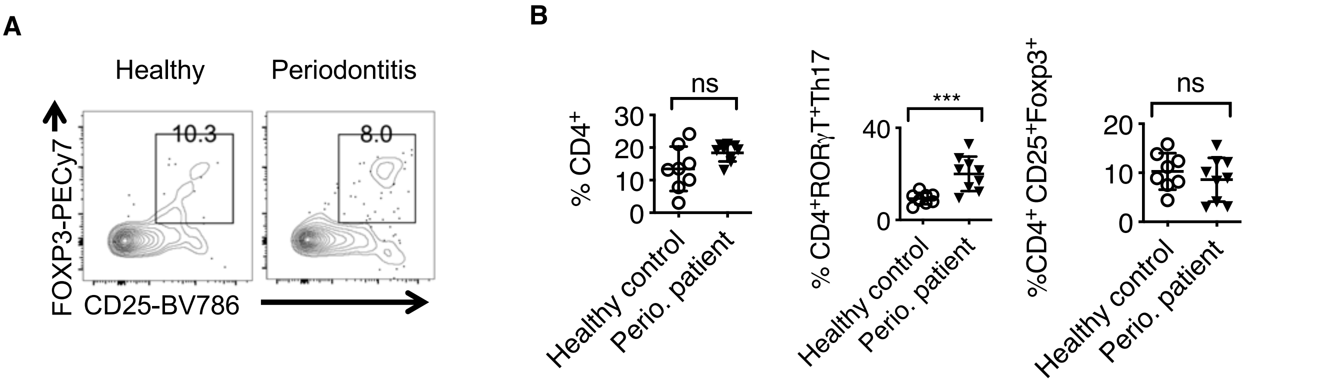


**Fig.S4. No $T_{reg}$ alterations were observed in the oral mucosa of periodontitis patients *ex vivo*.** HOILs from gingival mucosa of healthy control (healthy; n=8) or periodontitis patients (Perio. n= 9) were collected under a separate approved IRB protocol (UHCMC IRB number: 03-13-15). They were processed for flow cytometry. CD25 and FOXP3 expression in $CD3^+CD4^+$ gated HOIL cells (A). Statistical analyses comparing % CD4, Th17 cells and $T_{regs}$, comparing the two groups. * P< 0.05; Mann Whitney test.

Fig.S5

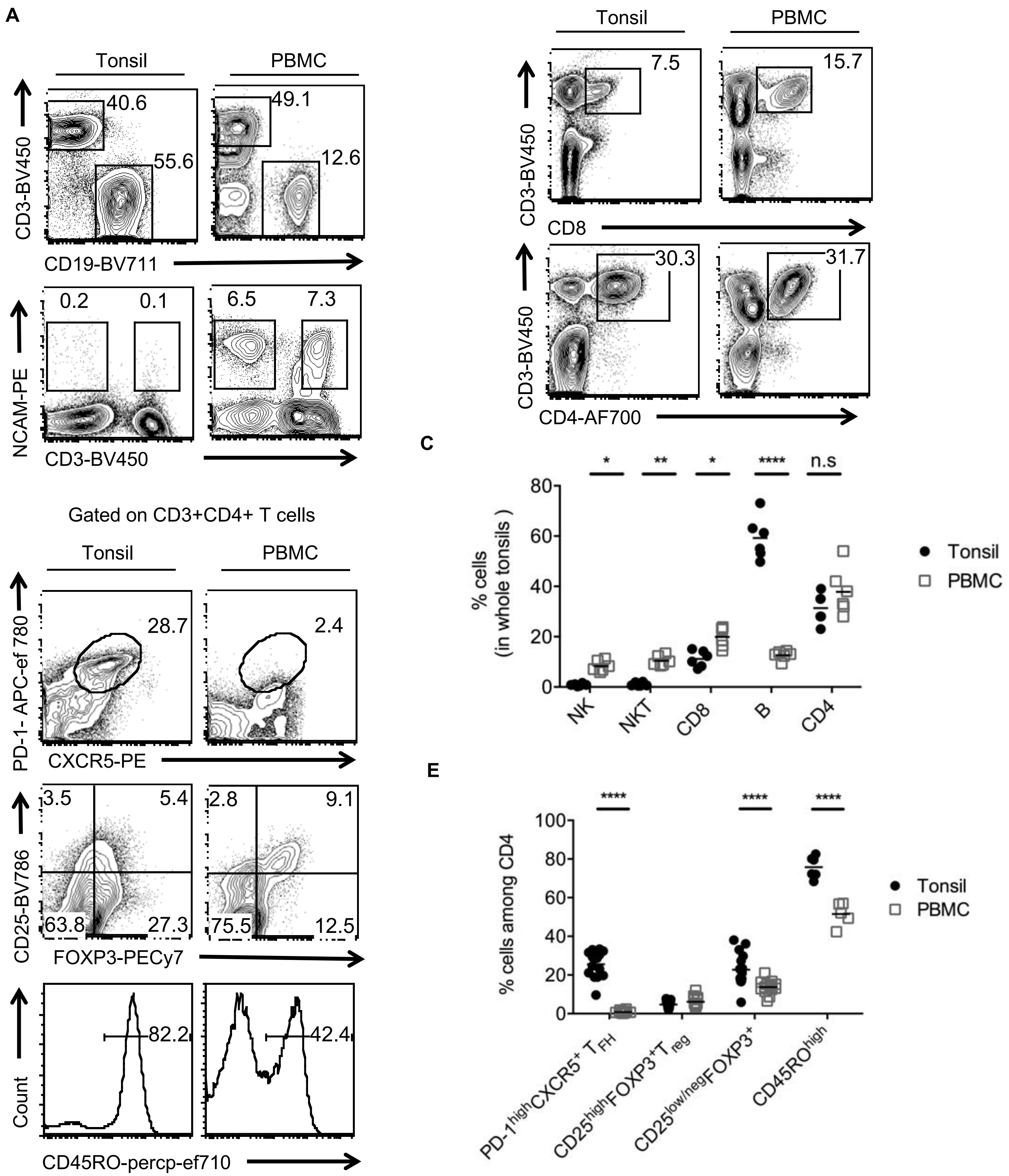


**Fig.S5. Characterization of immune cells in tonsils and comparison with PBMC cells *ex vivo*.** Tonsils and PBMC were processed *ex vivo*. (A) CD4 and CD8 cells in $CD3^+$ population (B), Statistical analyses and comparison between PBMC and tonsils for the indicated populations (C) and CXCR5, PD-1 (top), CD25, FOXP3 (middle) and CD45RO expression in $CD4^+$ T cells (D,E). Representative flow cytometric data and statistical analyses from 5 independent donors are shown.

Fig.S6

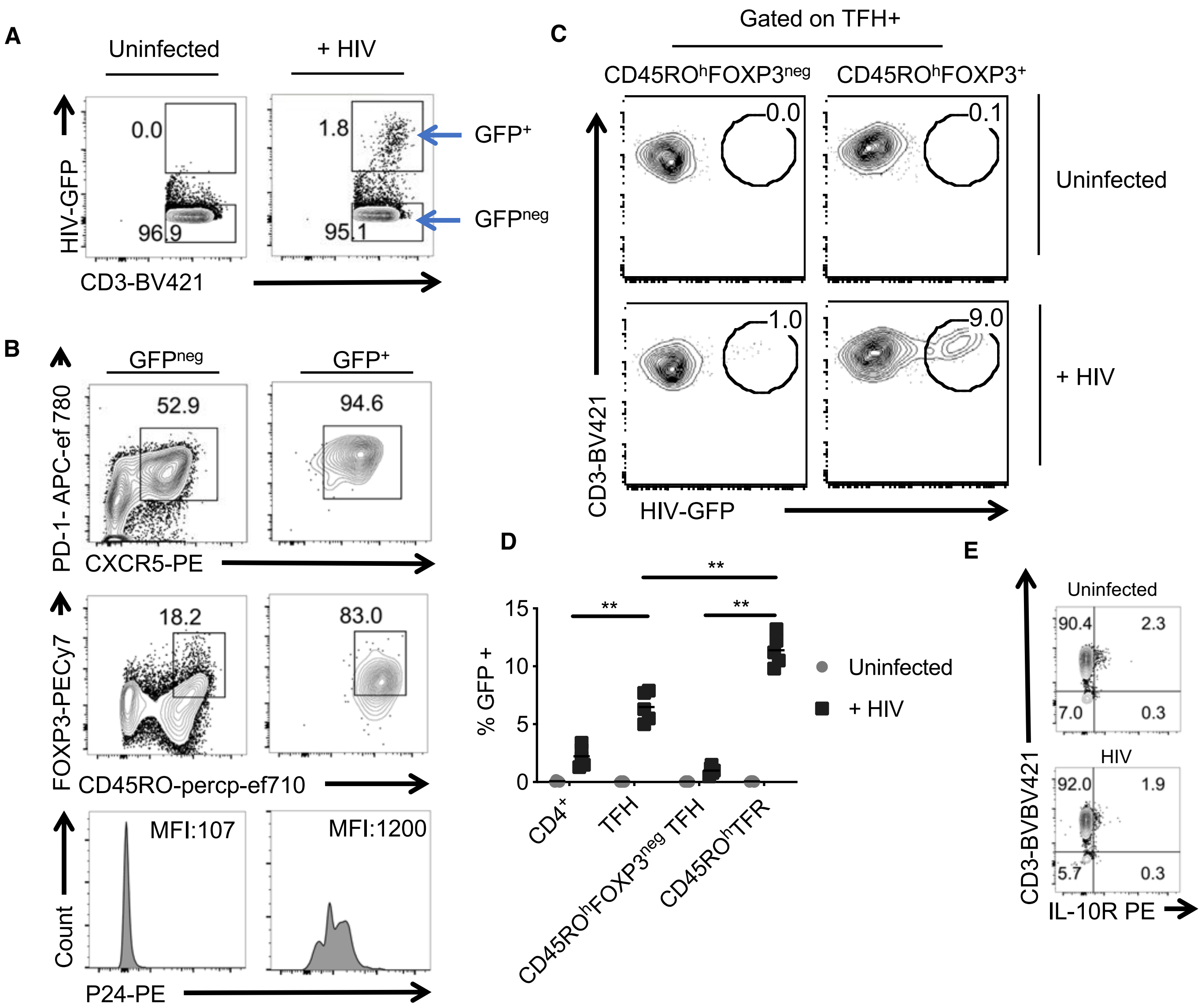


**Fig.S6. Characterization of HIV- infected cells in tonsils. A)** Whole human tonsil cultures (HTC) were activated by TCR stimulation and allowed to expand in the presence of TGF-β1 (10 ng/ml) and IL-2 (100 U/ml). They were infected with HIV on day 2 after TCR stimulation. Productive infection, as determined by GFP expression 72 hours post-infection in HTC. **B**) PD-1 and CXCR5 expressing TFH (top), CD45RO, FOXP3 (middle) and p24 expression in GFP negative and $GFP^{+}$ $CD4^{+}$ cells. **C**) GFP expression in $CD45RO^{high}FOXP3^{neg}$ and $CD45RO^{high}FOXP3^{+}$ cells in TFH population (above), statistical analyses of % GFP+ cells in indicated population (**D**). Representative flow cytometric data and statistical analyses from 5 independent tonsil donors are shown. (**E**) **IL-10 receptor was unaltered in $CD4^{+}$ cells in HIV-infected cultures** . Purified $CD4^{+}$ cells were activated and infected with HIV as above. Flow cytometry was performed to determine IL-10 receptor expression in $CD4^{+}$cells on 6 days post-infection. Representative flow cytometric data from two independent tonsil donors.

Fig.S7

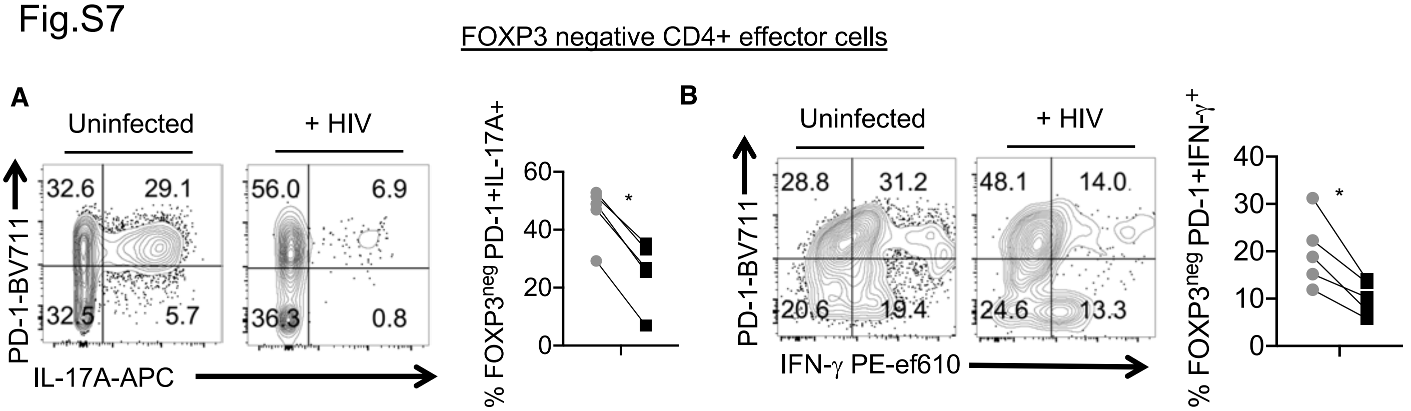


**Fig.S7. HIV- infection resulted in reduction of cytokine expressing CD4$^+$ T cell effectors in tonsils.** HTC were activated by TCR stimulation and infected as in Fig.3. PD-1 and IL-17A expression **A)**, and PD-1 and IFN-γ expression **B)** in FOXP3 negative CD4$^+$ effector cells 5 days post-infection.

Fig.S8

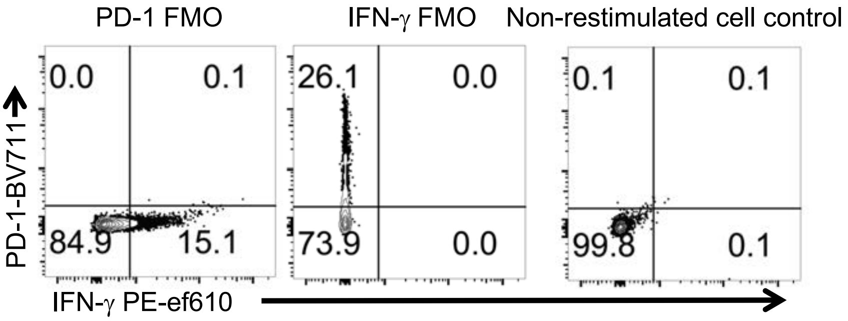


**Fig.S8. FMO controls.** HTC were activated were activated by TCR stimulation and allowed to expand in the presence of TGF-β1 (10 ng/ml) and IL-2 (100 U/ml). They were infected with HIV on day 2 after TCR stimulation. PD-1, IFN-γ FMO controls, and non-restimulated cell control for IFN-γ and PD-1 staining -gated on CD4$^+$FOXP3$^+$ cells 5 days post-infection.

Fig.S9

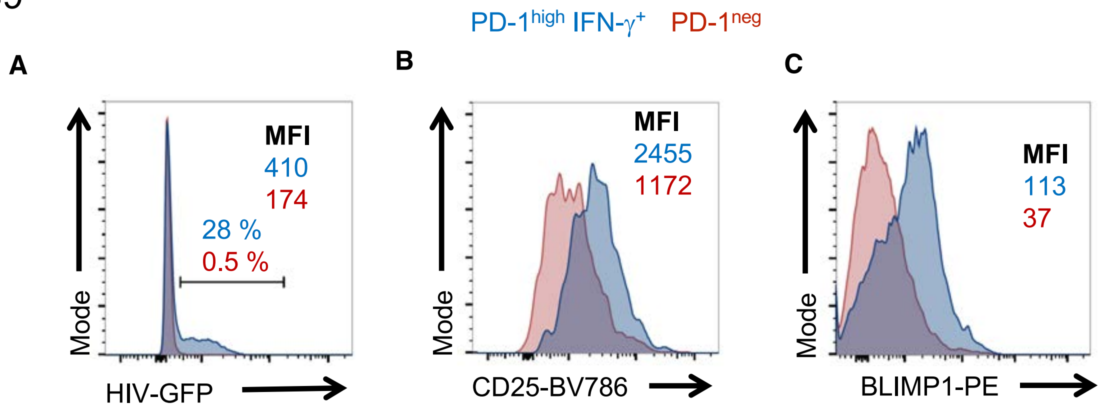


**Fig.S9. Characterization of PD-1$^{high}$ IFN-γ$^+$ and PD-1$^{low}$ cells in HIV- infected tonsils.** HTC were activated were activated by TCR stimulation and allowed to expand in the presence of TGF-β1 (10 ng/ml) and IL-2 (100 U/ml). They were infected with HIV on day 2 after TCR stimulation. GFP expression **A)** CD25 (**B**) and BLIMP-1 (**C**) expression in CD4$^+$FOXP3$^+$ cells 6 days post-infection. Representative flow cytometric data from 3 independent tonsil donors are shown.

Fig.S10

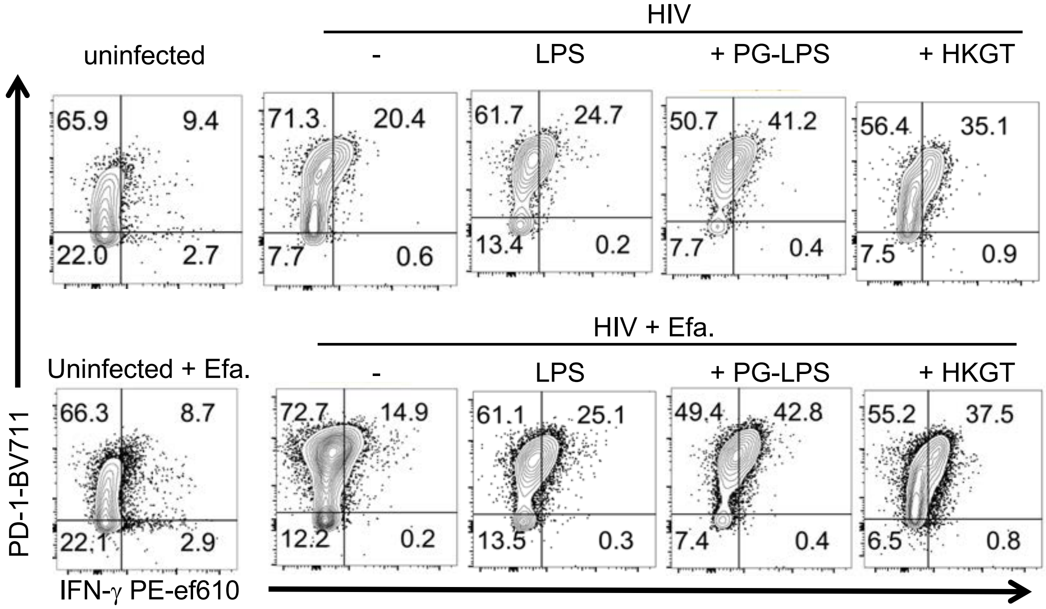


**Fig.S10. PD-1^hi^IFN-γ^+^ FOXP3^+^ cell accumulation is enhanced by TLR-2 ligands in the context of HIV infection. A)** Purified $CD4^+$ T cells were activated with TCR stimulation and allowed to expand in the presence of TGF-β1 (10 ng/ml) and IL-2 (100 U/ml). Some cells were infected with HIV on day 2 after TCR stimulation. Indicated cytokines or reagents were also added during this time. PD-1 and IFN-γ expression in $CD4^+FOXP3^+$ cells 6 days post-infection. Representative flow cytometric data from 3 independent tonsil donors are shown.

Fig.S11

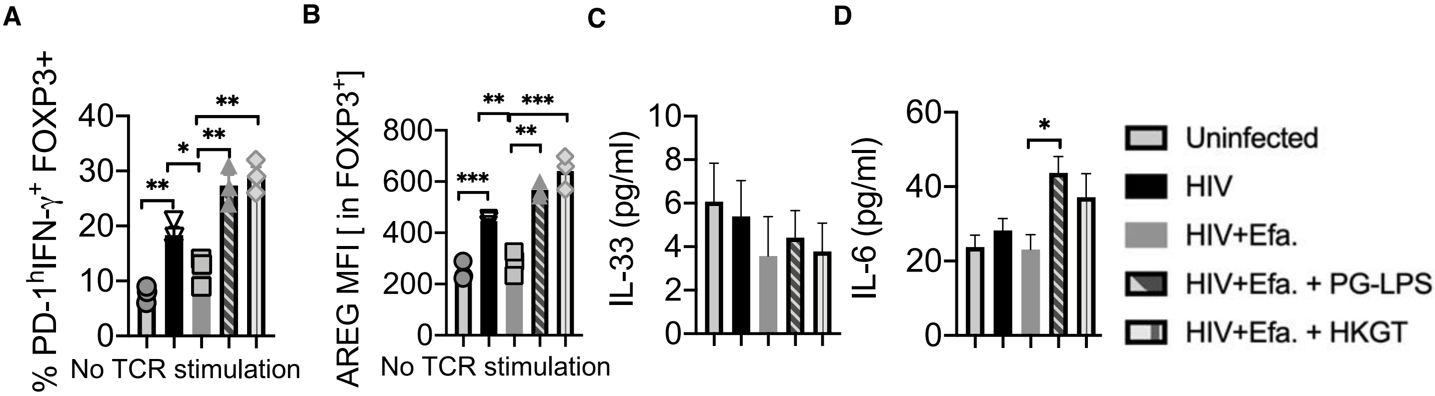


**Fig.S11. A,B) HIV induced PD-1^h^ IFN-γ^+^FOXP3^+^ and AREG expression in $FOXP3^+$ cells are independent of TCR stimulation.** Purified and unstimulated $CD4^+$ T cells allowed to expand in the presence of TGF-β1 (10 ng/ml) and IL-2 (100 U/ml). Some cells were infected with HIV on day 2 after TCR stimulation. Indicated cytokines or reagents were also added during this time. **C,D) Cytokine expression in tonsil $CD4^+$ T cells during HIV infection.** Purified $CD4^+$ T cells activated by TCR stimulation and infected as above. IL-33 (C) and IL-6 (D) were quantified by ELISA 3 days post-infection. Statistical analyses from 5 independent experiments.

Fig.S12

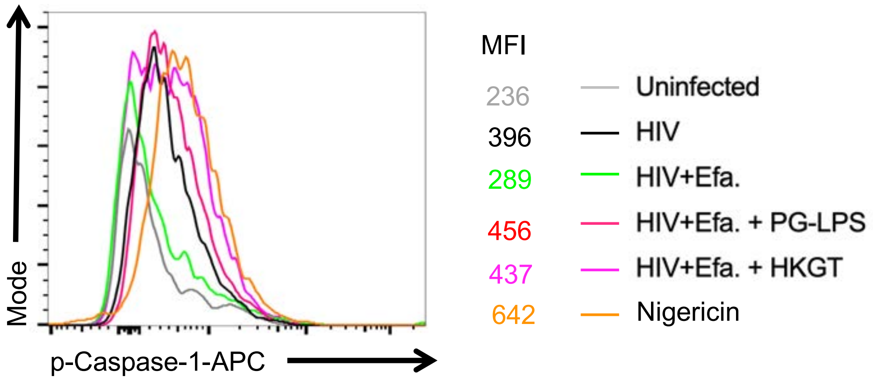


**Fig.S12. Activated caspase-1 expression in tonsil CD4$^+$ T cells during HIV infection.** Purified CD4$^+$ T cells were activated by TCR stimulation and allowed to expand in the presence of TGF-β1 (10 ng/ml) and IL-2 (100 U/ml). Some cells were infected with HIV on day 2 after TCR stimulation. Indicated cytokines or reagents were also added during this time. Phosphorylated caspase-1 expression 3 days post-infection. Representative flow cytometric data from 3 independent tonsil donors are shown.

Fig.S13

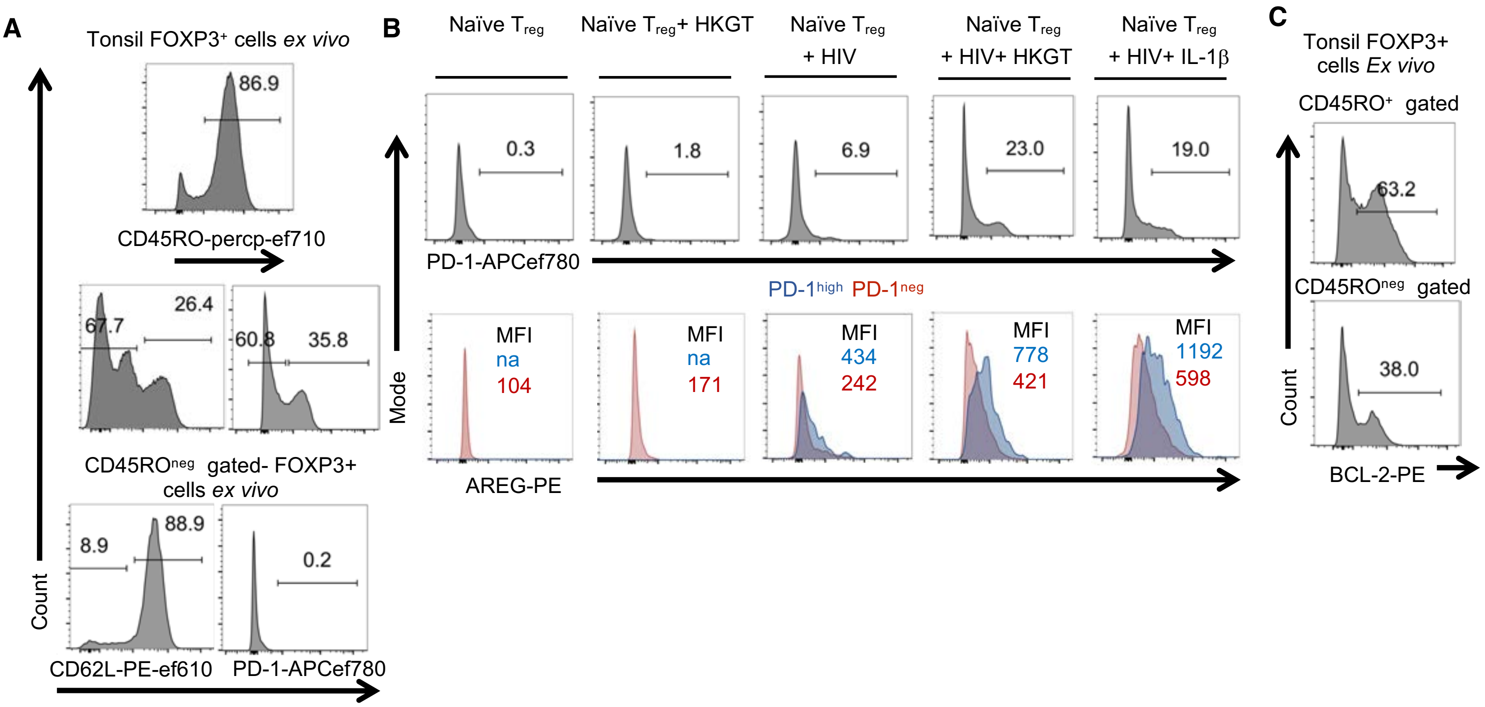


**Fig.S13. PD-1$^{hi}$IFN-γ$^+$ FOXP3$^+$ cell induction from naïve $T_{regs}$ is enhanced by TLR-2 ligands in the context of HIV infection.** A, C) *Ex vivo* flow cytometry analysis of tonsils (gated on CD4$^+$FOXP3$^+$ cells). B) CD4$^+$CD45RO$^{neg}$CD127$^{low}$CD25$^+$ $T_{reg}$ cells were sorted using three-step sorting of the tonsil cells. First, EasySep™ Human CD4+ T Cell Isolation Kit (STEMCELL Technologies) was used to isolate untouched CD4+ T cells using depletion of non-CD4 T cells. Then human CD45RO$^+$CD4$^+$ kit (Miltenyi biotech) was used to remove CD4 memory cells. Purified CD45RO$^{neg}$ naïve (92%) were used for further purification of $T_{regs}$ using human CD4+CD127lowCD25+ regulatory T cell kit (STEMCELL Technologies) (> 85% FOXP3$^+$). These cells were activated with TCR stimulation and allowed to expand in the presence of TGF-β1 (10 ng/ml) and IL-2 (100 U/ml). Some cells were infected with HIV on day 2 after TCR stimulation. Indicated cytokines or reagents were also added during this time. Flow cytometry was performed on day 7 after infection. Representative flow cytometric data from 3 independent tonsil donors are shown.

Fig.S14

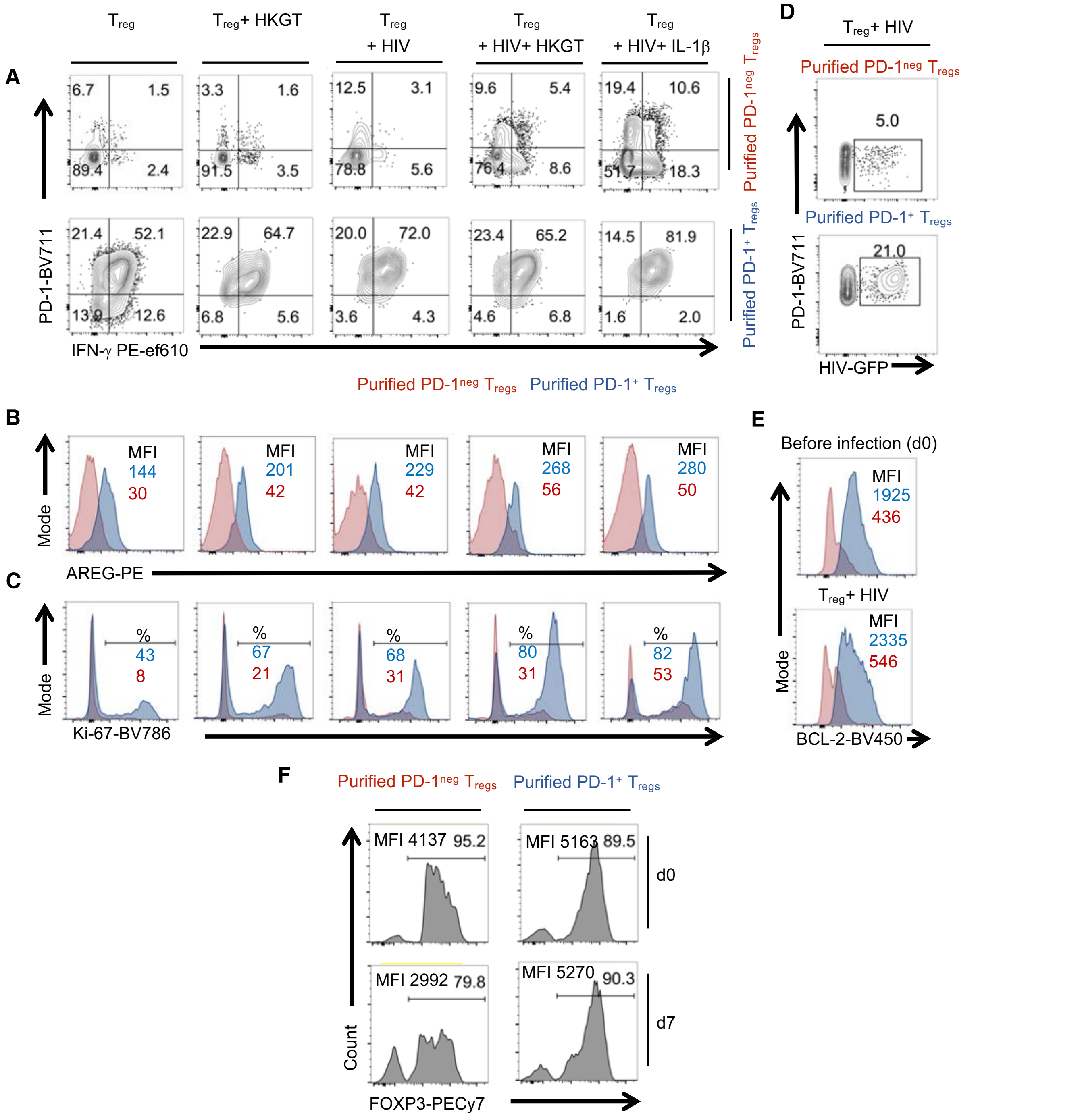


**Fig.S14. PD-1$^{hi}$IFN-γ$^{+}$ FOXP3$^{+}$ cell induction and proliferation are enhanced by TLR-2 ligands in the context of HIV infection.** CD4$^{+}$PD-1$^{neg}$CD127$^{low}$CD25$^{+}$ T$_{reg}$ and CD4$^{+}$PD-1$^{+}$CD127$^{low}$CD25$^{+}$ T$_{reg}$ cells were sorted using three-step sorting of the tonsil cells. EasySep™ Human CD4+ T Cell Isolation Kit (STEMCELL Technologies) was used to isolate untouched CD4$^{+}$ T cells using depletion of non-CD4 T cells. Anti-Biotin multisort kit (Miltenyi biotech) was used to positively sort anti-PD-1-biotin labelled cells. The beads were removed from positively sorted cells. Purified CD4$^{+}$PD-1$^{+}$ and CD4$^{+}$PD-1$^{neg}$ (~90-95%) were used for further purification of T$_{regs}$ using human CD4+CD127lowCD25+ regulatory T cell kit (STEMCELL Technologies) (> 85% FOXP3$^{+}$). These cells were activated with TCR stimulation and allowed to expand in the presence of TGF-β1 (10 ng/ml) and IL-2 (100 U/ml). Some cells were infected with HIV on day 2 after TCR stimulation. Indicated cytokines or reagents were also added during this time. Flow cytometry was performed on day 7 after infection to determine PD-1, IFN-γ, (A), AREG (B), Ki-67 (C), and GFP (D). BCL-2 (E) and FOXP3 (F) expression on day 0 (d0) and d7 after HIV infection. Representative flow cytometric data from 3 independent tonsil donors are shown.

Fig.S15

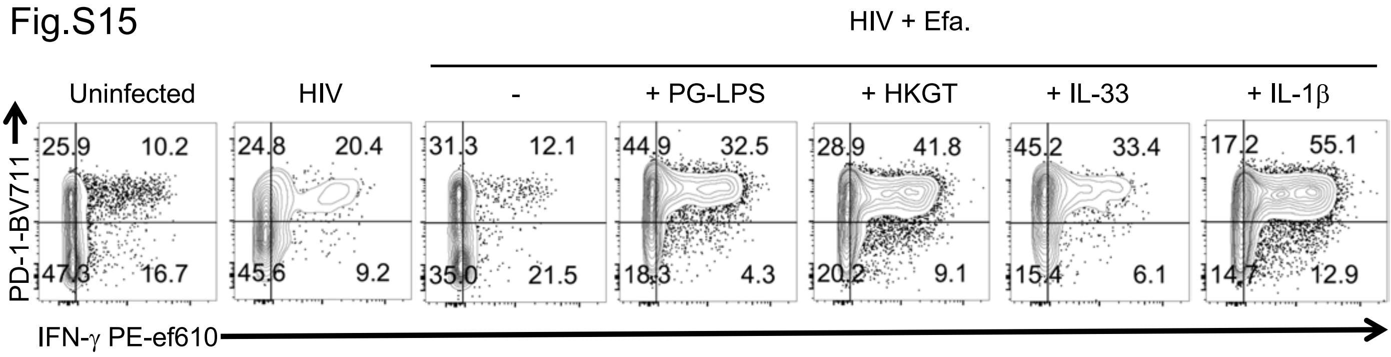


**Fig.S13. PD-1$^{hi}$IFN-γ$^{+}$ FOXP3$^{+}$ cell induction is enhanced by TLR-2 ligands, IL-33 and IL-1β in the context of HIV infection.** Purified CD4$^{+}$ T cells were activated by TCR stimulation and allowed to expand in the presence of TGF-β1 (10 ng/ml) and IL-2 (100 U/ml). Some cells were infected with HIV on day 2 after TCR stimulation. Indicated cytokines or reagents were also added during this time. PD-1 and IFN-γ expression in CD4$^{+}$FOXP3$^{+}$ cells 6 days post-infection.

Fig.S16

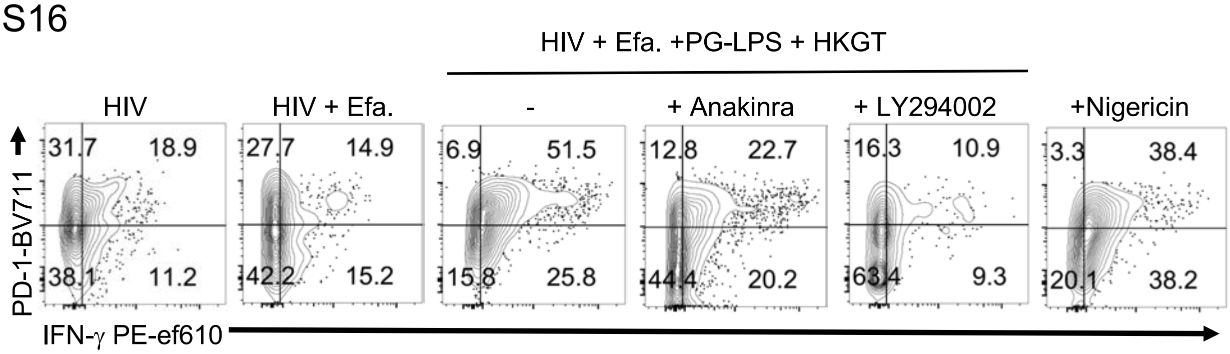


**Fig.S16. PD-1$^{hi}$IFN-γ$^{+}$ FOXP3$^{+}$ cell induction requires Akt-1 dependent IL-1β and inflammasome signaling in the context of HIV infection.** Purified CD4$^{+}$ T cells were were activated by TCR stimulation and allowed to expand in the presence of TGF-β1 (10 ng/ml) and IL-2 (100 U/ml). Some cells were infected with HIV on day 2 after TCR stimulation. Indicated cytokines or reagents were also added during this time. PD-1 and IFN-γ expression in CD4$^{+}$FOXP3$^{+}$ cells 6 days post-infection. Representative flow cytometric data from five independent tonsil donors are shown.

Fig.S17

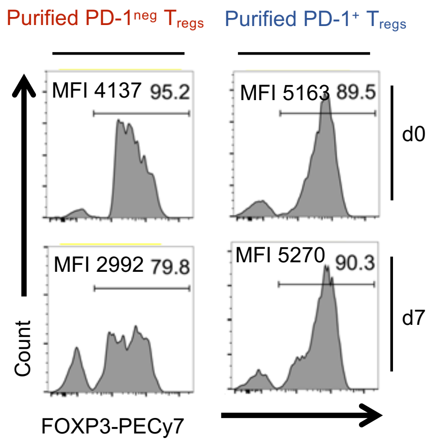


**Fig.S18. PD-1$^{neg}$ FOXP3$^{+}$ cells lose FOXP3 expression in the context of HIV infection.** CD4$^{+}$PD-1$^{neg}$CD127$^{low}$CD25$^{+}$ $T_{reg}$ and CD4$^{+}$PD-1$^{+}$CD127$^{low}$CD25$^{+}$ $T_{reg}$ cells were sorted using three-step sorting of the tonsil cells. EasySep™ Human CD4+ T Cell Isolation Kit (STEMCELL Technologies) was used to isolate untouched CD4$^{+}$ T cells using depletion of non-CD4 T cells. Anti-Biotin multisort kit (Miltenyi biotech) was used to positively sort anti-PD-1-biotin labelled cells. The beads were removed from positively sorted cells. Purified CD4$^{+}$PD-1$^{+}$ and CD4$^{+}$PD-1$^{neg}$ (~90-95%) were used for further purification of $T_{regs}$ using human CD4+CD127lowCD25+ regulatory T cell kit (STEMCELL Technologies) (> 85% FOXP3$^{+}$). These cells were activated with TCR stimulation and allowed to expand in the presence of TGF-β1 (10 ng/ml) and IL-2 (100 U/ml). Some cells were infected with HIV on day 2 after TCR stimulation. Flow cytometry was performed to determine FOXP3 expression on day 0 (d0) and d7 after HIV infection. Representative flow cytometric data from 3 independent tonsil donors are shown.

Fig.S18

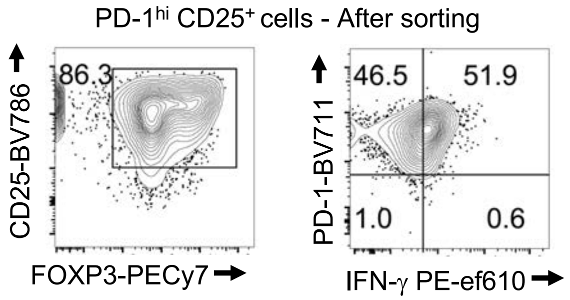


**Fig.S18. PD-1$^{hi}$CD25$^{+}$ cell purity.** Purified CD4$^{+}$ T cells were activated with TCR stimulation and allowed to expand in the presence of TGF-β1 (10 ng/ml) and IL-2 (100 U/ml). Some cells were infected with HIV on day 2 after TCR stimulation. . PD-1$^{hi}$CD25$^{+}$ cells were purified from HIV-infected CD4 cultures using sequential sorting of PD-1-PE$^{+}$ cells and CD25$^{high}$ $T_{reg}$ cells using STEMCELL technology PE isolation and CD25+Treg isolation kits. Data represent three independent experiments.

Fig.S19

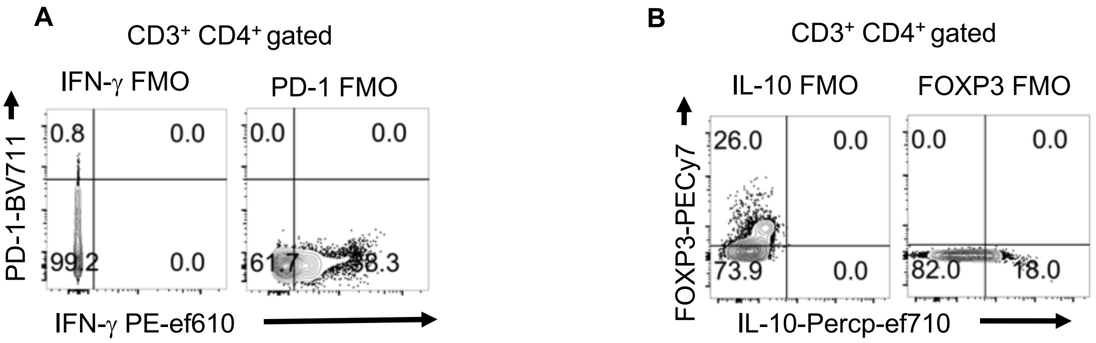


**Fig.S19. FMO controls.** PD-1 and IFN-γ (A) IL-10 and FOXP3 FMO controls gating on CD4$^{+}$ cells in gingival mucosa processed for flow cytometry *ex vivo*. Cells were restimulated with PMA/Ionomycin for 4 hours before flow cytometry. The gates were assigned based on the unstained controls, FMO controls, and non-restimulated control cells, for each experiment. Because tissue cells (oral mucosa) may have some autofluorescence, we also performed PBMC staining control cells in parallel. We chose the most appropriate gating based on an objective evaluation using these controls when establishing the staining protocol for experiments. Representative flow cytometric data from 10-12 independent study participants are shown.
